# Dissociable encoding of motivated behavior by parallel thalamo-striatal projections

**DOI:** 10.1101/2023.07.07.548113

**Authors:** Sofia Beas, Isbah Khan, Claire Gao, Gabriel Loewinger, Emma Macdonald, Alison Bashford, Shakira Rodriguez-Gonzalez, Francisco Pereira, Mario A. Penzo

## Abstract

The successful pursuit of goals requires the coordinated execution and termination of actions that lead to positive outcomes. This process is thought to rely on motivational states that are guided by internal drivers, such as hunger or fear. However, the mechanisms by which the brain tracks motivational states to shape instrumental actions are not fully understood. The paraventricular nucleus of the thalamus (PVT) is a midline thalamic nucleus that shapes motivated behaviors via its projections to the nucleus accumbens (NAc)^1–8^ and monitors internal state via interoceptive inputs from the hypothalamus and brainstem^3,9–14^. Recent studies indicate that the PVT can be subdivided into two major neuronal subpopulations, namely PVT^D2(+)^ and PVT^D2(–)^, which differ in genetic identity, functionality, and anatomical connectivity to other brain regions, including the NAc^4,15,16^. In this study, we used fiber photometry to investigate the *in vivo* dynamics of these two distinct PVT neuronal types in mice performing a reward foraging-like behavioral task. We discovered that PVT^D2(+)^ and PVT^D2(−)^ neurons encode the execution and termination of goal-oriented actions, respectively. Furthermore, activity in the PVT^D2(+)^ neuronal population mirrored motivation parameters such as vigor and satiety. Similarly, PVT^D2(−)^ neurons, also mirrored some of these parameters but to a much lesser extent. Importantly, these features were largely preserved when activity in PVT projections to the NAc was selectively assessed. Collectively, our results highlight the existence of two parallel thalamo-striatal projections that participate in the dynamic regulation of goal pursuits and provide insight into the mechanisms by which the brain tracks motivational states to shape instrumental actions.

## RESULTS

### Characterization of motivated behavior in rodents using a foraging-like task

Goal-oriented behaviors are generated by cortico-mesolimbic circuits that establish goal objects based on need states, as well as the actions required to obtain those objects^17^. Recent reports suggest that the PVT plays a critical role in translating needs states into motivation, owing to its strong innervation by the hypothalamus and brainstem^10,11,13,14,18^. The PVT is composed of molecularly diverse neuronal subpopulations that segregate across the anteroposterior axis of the thalamus and project to the NAc^8,15,16,19^. Particularly, the PVT contains two major distinct subpopulations, termed PVT^D2(+)^ and PVT^D2(−)^, respectively, identified based on the expression of the dopamine D2 receptor, or lack thereof^4,16^. However, how activity within parallel thalamo-striatal projections arising in the PVT relates to specific aspects of motivated behaviors remains unclear.

To address this question, we investigated the *in vivo* dynamics of PVT^D2(+)^ and PVT^D2(−)^ neurons using a reward foraging-like behavioral task^11,20^. In this task, mice initiated individual trials by entering the “trigger zone” where they were presented with a cue that signaled reward availability. Upon cue presentation, mice were required to traverse a long corridor and reach the “reward zone” to obtain a food reward (Supp. Fig. 1A) (See Methods). We found that mice successfully learned to shuttle back and forth to obtain the food reward and were highly engaged in the task, as demonstrated by two observations. First, they completed a significantly larger volume of trials in late compared to early sessions (Supp. Fig. 1B). Second, they displayed shorter latencies to reach the reward zone (Supp. Fig. 1C), obtain the reward (Supp. Fig. 1D), and initiate subsequent trials (Supp. Fig. 1E).

We noticed that mice varied in latencies to perform individual trials, but after training, they completed most trials under 11 seconds (∼ 75% of completed trials; Supp. Fig. 1F). Thus, trial engagement was denoted by trials completed in 11 seconds or less. As such, to investigate how molecularly distinct subpopulations of the PVT encode various parameters associated with goal pursuit, the analysis of calcium signals recorded via fiber photometry was restricted to trials meeting the engagement criterion (Supp. Fig. 1F). Importantly, this trial inclusion criteria allowed us to have a range of trials with different reward zone and reward delivery latencies that did not significantly differ throughout testing sessions (Supp. Fig. 1F) (See Methods). Moreover, it provided a suitable time frame for investigating the activity of PVT^D2(+)^ and PVT^D2(−)^ neurons during different trial groups and during different stages of the trials.

### pPVT^D2(+)^ neuronal activity dynamics during food-seeking behavior

We measured the activity of posterior PVT^D2(+)^ (pPVT^D2(+)^) neurons while food-restricted mice performed trials in our foraging-like reward task. For this, we injected the PVT of *Drd2*-Cre mice with an AAV driving expression of a Cre-dependent calcium sensor (GCaMP6s) and recorded bulk fluorescence signals from this neuronal population using fiber photometry (See Methods; Fig. 1A). At cue onset, we found no changes in the averaged calcium signal in this population (Supp. Fig. 2A). However, we did find that pPVT^D2(+)^ neurons were robustly activated as mice approached the reward zone and the receptacle to obtain the food reward (strawberry Ensure^®^; Fig. 1B). Modest but statistically significant increases in calcium signals were also observed in these neuronal populations upon reward delivery (Fig. 1C). Importantly, no such changes in fluorescence were observed in control mice in which pPVT^D2(+)^ neurons expressed GFP instead of GCaMP (Fig. 1D-F). Collectively, these results demonstrate that pPVT^D2(+)^ neurons are modulated by reward-seeking and consumption.

**Figure 1.**
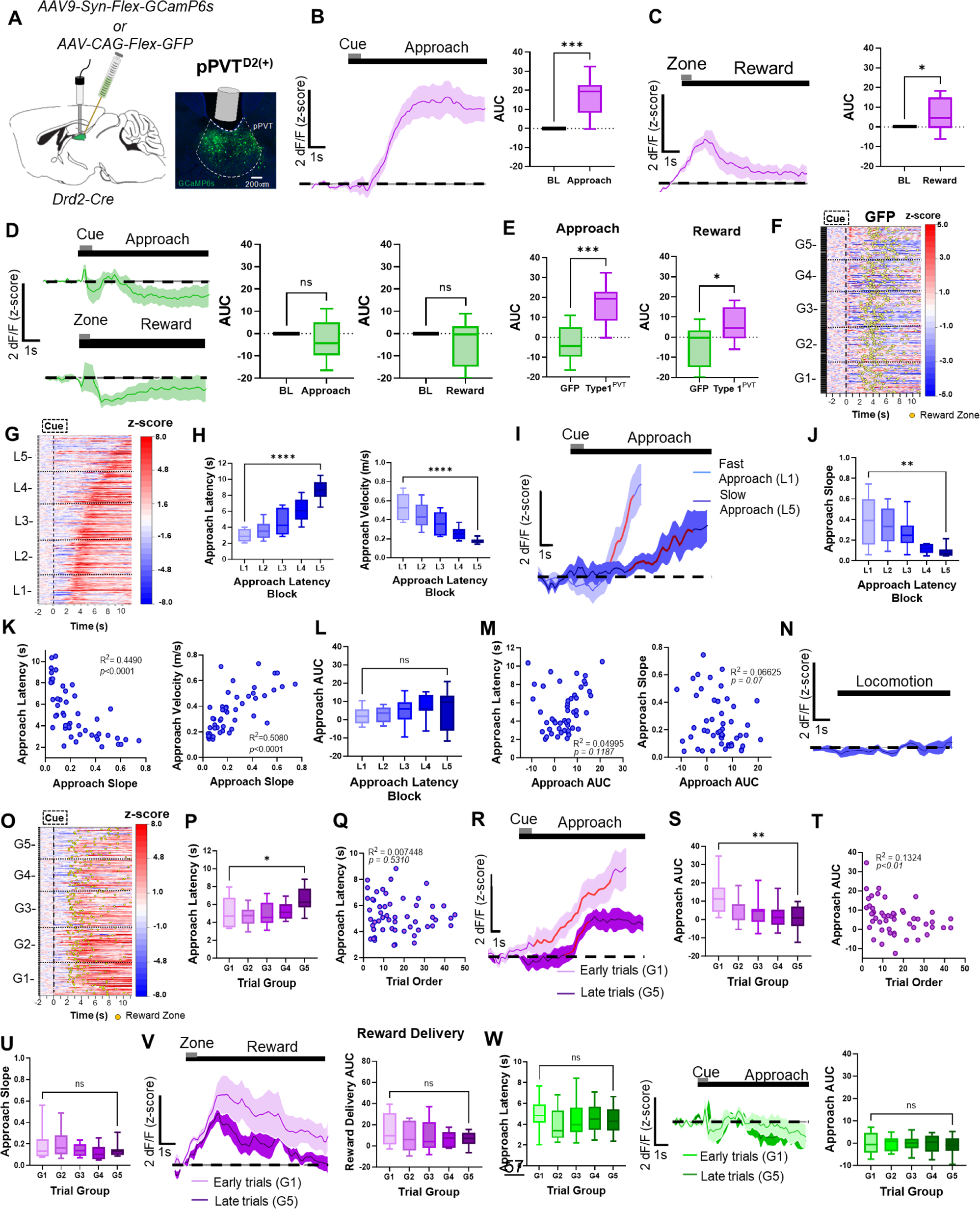
*In-vivo* activity dynamics of PVT^D2(+)^ neurons. **(A)** *Left*: Schematic of the stereotaxic injections and optical fiber implantation for targeting D2(+) neurons in the pPVT. *Right*: Representative image from a Drd2-Cre mouse expressing GCaMP6s in the pPVT. **(B)** *Left*: Average GCaMP6s responses from pPVT^D2(+)^ neurons showing ramping activity during reward approach. *Right*: Quantification of the approach-evoked changes in GCaMP6s fluorescence in pPVT^D2(+)^ neurons. Area under the curve (AUC), n = 296 trials from 6 mice, two-tailed paired t-test, *\*\*\*p<0.001*. **(C)** *Left*: Average GCaMP6s responses from pPVT^D2(+)^ neurons during reward delivery. *Right*: AUC quantification of the reward-evoked changes in GCaMP6s fluorescence in pPVT^D2(+)^ neurons. n = 296 trials from 6 mice, two-tailed paired t-test, *\*p<0.05*. **(D)** *Left*: Top *-* Average GFP fluorescence during reward approach. Bottom -Average GFP activity during reward delivery. *Middle:* AUC quantification of GFP fluorescence during the baseline and approach periods. Two-tailed paired t-test, *p=0.29*; ns, not significant. *Right:* AUC quantification of GFP fluorescence during the baseline and reward delivery periods. Two-tailed paired t-test, *p=0.18*; ns, not significant. **(E)** *Left*: AUC quantification between GFP and pPVT^D2(+)^ neuronal activity during reward approach. Two-tailed unpaired t-test, n = 544 trials from 5 mice, *\*\*\*p<0.001. Right:* AUC quantification between GFP and pPVT^D2(+)^ neuronal activity during reward delivery. AUC, Two-tailed unpaired t-test, *\*p<0.05*. **(F)** Heatmap of GFP activity in the pPVT, time-locked to cue onset, sorted by trial order, and binned into 5 ‘trial group blocks’ (G1 – G5). Yellow dots represent reward zone arrival. **(G)** Heatmap showing excitatory reward approach responses from pPVT^D2(+)^ neurons, time-locked to cue onset, sorted by latency to approach the reward zone and binned into 5 ‘approach latency blocks,’ L1, n = 61 trials, L2, n= 67 trials, L3, n = 67 trials, L4, n = 67 trials, L5, n = 64 trials from 6 mice. **(H)** *Left*: Latencies to reach the reward zone in seconds for each approach latency block. Repeated measures ANOVA, *\*\*\*\*p<0.0001. Right*: Velocity (m/s) during reward approach calculated for each approach latency block. Repeated measures ANOVA, *\*\*\*\*p<0.0001*. **(I)** Average GCaMP6s responses for fast and slow reward approach. The red line represents 20-80% of the slope of the line. **(J)** Slope-of-the-line quantifications of GCaMP6s activity from pPVT^D2(+)^ neurons across approach latency blocks. Repeated measures ANOVA, *\*\*p<0.01*. **(K)** *Left:* Pairwise correlation between the approach latency and the slope-of-the-line quantifications of GCaMP6s responses from pPVT^D2(+)^ neurons during the approach. *Right:* Correlation between the velocity (m/s) during the approach and the slope-of-the-line quantifications of GCaMP6s responses from pPVT^D2(+)^ neurons during the approach. Each dot represents the GCaMP6s activity of individual mice averaged across approach latency blocks. **(L)** AUC quantifications of the reward approach-evoked changes in GCaMP6s activity across approach latency blocks. Repeated measures ANOVA, *p=0.55;* ns, not significant **(M)***. Left:* No correlation between approach latency and AUC. *Right:* No correlation between the AUC and the slope-of-the-line quantifications of GCaMP6s responses from pPVT^D2(+)^ neurons during the approach. **(N)** Average GCaMP6s fluorescence from pPVT^D2(+)^ neurons during spontaneous locomotion. **(O)** Same as (G), but responses were sorted by trial order and binned into 5 ‘trial group blocks.’ G1, n = 61 trials, G2, n= 67 trials, G3, n = 67 trials, G4, n = 67 trials, G5, n = 64 trials from 6 mice. Yellow dots represent reward zone arrival. **(P)** Latencies to reach the reward zone across trial group blocks. Repeated measures ANOVA, *\*\*p<0.001*. **(Q)** No correlation between the approach latency and trial order. **(R)** Average GCaMP6s responses from pPVT^D2(+)^ neurons comparing approach trials performed early and late in the testing session. The red line indicates 20-80% of the slope of the line. **(S)** AUC quantification of the reward approach-evoked changes in GCaMP6s activity across trial group blocks. Repeated measures ANOVA, *\*\*p<0.001*. **(T)** Correlation between the AUC of the reward approach-evoked changes in GCaMP6s activity and trial order within a test session. **(U)** Slope-of-the-line quantifications of GCaMP6s activity across trial group blocks. Repeated measures ANOVA, *p=0.36*. **(V)** *Left:* Average GCaMP6s responses from pPVT^D2(+)^ neurons upon reward delivery comparing early and late trials in the testing session. *Right:* AUC quantification during reward delivery across trial group blocks. Repeated measures ANOVA, *p=0.36*; ns, not significant. **(W)** *Left:* Latencies to reach the reward zone across trial group blocks. Repeated measures ANOVA, *p=0.25;* ns, not significant. *Middle:* Average GFP activity in the pPVT comparing approach trials performed early or late in the testing session. *Right:* AUC quantification of GFP activity in the pPVT during approach across trial group blocks. Repeated measures ANOVA*, p=0.60*; ns, not significant. All data in the figure are shown as mean ±s.e.m.

To determine whether the activity of pPVT^D2(+)^ neurons varied with the motivational state of the subjects, we ranked all trials based on the latency to reach the reward zone (Fig. 1G) and divided them into five categories (L1–L5; Supp. Fig. 1G). Trials with the shortest latencies to the reward zone (i.e., trials performed within 2-3 sec; Fig. 1H) were categorized as ‘fast trials’ (L1), whereas trials with the longest latencies (i.e., trials performed within 9-11 sec; Fig. 1H) were categorized as ‘slow trials’ (L5). The trial distribution for these two categories is shown in Supplemental Figure 1I, J. We next compared the activity of pPVT^D2(+)^ neurons during fast trials vs. slow trials. First our analyses showed that the cue presentation calcium signal did not differ between fast or slow approaches (Supp. Fig. 2B). However, during fast approach, pPVT^D2(+)^ neurons appeared to display higher GCaMP signal ramps compared to slow approach (Fig. 1I). These differences in the GCaMP signal ramps were quantified by calculating the 20-80% slope of the peak of GCaMP transients observed during the reward approach. Quantifications of the signal’s slope confirmed statistically significant differences in the activity of pPVT^D2(+)^ neurons between fast and slow trials (Fig. 1J). Moreover, we found that both latency and velocity parameters were negatively correlated with the GCaMP signals (Fig. 1K). Surprisingly, quantifications of the area under the curve (AUC) of the signal (Fig.1L) revealed no significant differences between fast and slow trials. Accordingly, no association was found between the latency to complete trials and signal AUC or between the approach AUC and the approach slope (Fig. 1M). Lastly, no changes in the activity of pPVT^D2(+)^ neurons were observed when mice were moving around the maze but were not engaged in the task (Fig. 1N). Altogether, our findings thus far suggest that pPVT^D2(+)^ neurons are modulated by reward-seeking and consumption and that their activity varies with motivation.

To test if changes in need states impact pPVT^D2(+)^ neuronal activity, we sorted trials by trial order, categorizing the earliest trials as ‘G1’ and the latest trials as ‘G5.’ (Fig. 1O - U). We implemented this strategy to model a satiety component, which assumes that well-trained food-restricted mice are hungrier at the beginning compared to the end of each 60 min testing session^21^. Importantly, the average latency to approach reward did not show statistically significant differences across five trial groups distributed throughout testing sessions (Supp. Fig. 1F; Fig.1Q). Previous research has shown that NAc-projecting neurons of the pPVT are engaged by hunger signals and changes in glucose levels, suggesting that need states influence the activity of PVT neurons^11,13,22^. In agreement with this view, we found that the activity of pPVT^D2(+)^ neurons was higher during early compared to later trials, indicating that pPVT^D2(+)^ neuronal activity is modulated by the satiety levels of subjects (Fig. 1R-T). Of note, quantifications of the slope revealed no significant differences between early and late trials (Fig. 1U). We also found that the activity of pPVT^D2(+)^ neurons at reward delivery (Fig. 1V) and during cue presentation (Supp. Fig 2C) trended towards a reduction as the session progressed, but these changes were not statistically significant. Importantly, the observed decreases in the activity of pPVT^D2(+)^ neurons were unlikely due to photobleaching of the GCaMP signal, as no statistically significant decreases in fluorescence signal were observed in GFP controls (Fig.1W). To probe our conclusions further, GCaMP-expressing mice in pPVT^D2(+)^ neurons were tested in a separate session where the reward was completely omitted for all trials. We observed that, under these conditions, these neurons were highly active across trials (Supp. Fig. 2D, E), and unlike for rewarded trials, this activity did not vary throughout sessions (Supp. Fig. 2F,G). Notably, ‘fast trials’ still displayed significantly higher GCaMP ramps compared to ‘slow trials’, again supporting the notion that the activity dynamics of pPVT^D2(+)^ neurons reflect behavioral vigor (Supp. Fig. 2H, I). Altogether, these findings suggest that pPVT^D2(+)^ neuronal activity is influenced by satiety levels, with higher activity observed early on in test sessions and lower activity observed towards the end of sessions as satiety levels increase.

### Functional linear mixed model analysis of trial-level temporal dynamics for pPVT^D2(+)^ neuronal activity

Up to this point, we performed analyses of photometric signals for pPVT^D2(+)^ neurons based on either latency or trial order. However, these analyses involved condensing trial types into summary measures, which obscures trial-level information. To mitigate this limitation, we employed a new statistical framework for the analysis of fiber photometry signals based on functional linear mixed modeling (FLMM)^23,24^ to further explore the relationship between the temporal structure of photometric signals and task-related covariates. FLMM combines mixed-effects modeling and functional regression to account for nested experimental studies containing multiple events in a trial and autocorrelations in the photometry signal (see Methods). The result of this analysis is a single plot of the coefficient estimates for each covariate (e.g., approach latency), which shows the timepoints at which covariates have a statistically significant association with the photometry signal across trials. Using FLMM, we then tested the effects of latency and trial order during the following approach-related events: 1. cue presentation; and 2. reward zone arrival (Supp. Fig. 3). For pPVT^D2(+)^ neurons, FLMM showed an effect of trial order but not latency on the magnitude of the photometry signal (Supp. Fig. 3A, B). The lack of latency effect on signal is surprising considering that GCaMP signal in pPVT^D2(+)^ neurons rose more rapidly for fast compared to slow trials (Fig. 1J). One potential reason for this discrepancy is that FLMM was performed on relatively short time series surrounding discrete events. As such, when aligning signal to cue onset and reward zone arrival, a significant fraction of approach-related signal dynamic might have been missed. Of note, FLMM analysis generally yielded more nuanced information of how photometry signal collected from pPVT^D2(+)^ relates to behavior. Specifically, we found a trial order effect on pPVT^D2(+)^ neuronal activity during cue presentation and during reward zone entry. These effects revealed a negative relationship between trial order and photometry signal, such that as the mice performed more trials in a session, the pPVT^D2+^ neuronal signal significantly decreased during cue presentation and around reward zone entry (Supp Fig. 3B). Altogether, FLMM offers a robust statistical framework that can extract meaningful information about the temporal structure of calcium traces across trials and subjects in a task. As such, for the remainder of our study, we contrast our observations based on summary measures with those yielded by FLMM.

### PVT^D2(−)^ neuronal activity dynamics during food-seeking behavior

We next examined the PVT^D2(−)^ neuronal population and asked whether their activity dynamics during the reward foraging task differ from that of PVT^D2(+)^ neurons. For this, the pPVT of *Drd2*-Cre mice was injected with an AAV driving expression of GCaMP6s exclusively in Cre negative neurons (CreOFF; pPVT^D2(−)^) and fiber photometry was used to measure their activity in well-trained mice performing the reward task (Fig. 2A). We found that, unlike pPVT^D2(+)^ neurons, the activity of pPVT^D2(−)^ neurons significantly decreased during both reward approach and reward delivery (Fig. 2B, C). Given that, like pPVT^D2(+)^ neurons, the activity of pPVT^D2(−)^ neurons is modulated during reward approach and delivery, we then sought to assess whether the activity of pPVT^D2(−)^ neurons could also be influenced by motivational factors. For this, we grouped trials based on their latency to reach the reward zone as described earlier (Fig. 2D, E; Supp. Fig. 1H, K, L). Interestingly, the slope of downward ramps did not significantly differ between the two trial types (Fig. 2F, G), nor was there any correlation between the slope and the latency to reward approach or trial velocity (Fig. 2H, I). There was no statistically significant effect on AUC for fast and slow trials (Fig. 2J). These findings indicate that pPVT^D2(−)^ neuronal activity decreases during the reward approach, but these decreases might not be associated with motivation-related parameters such as latency to complete trials. Next, to investigate whether need states impact the activity of pPVT^D2(−)^ neurons, we categorized trials by trial order as noted earlier (Fig. 2K, L). Our analyses revealed that unlike for pPVT^D2(+)^ neurons, GCaMP signal recorded from pPVT^D2(−)^ neurons displayed no statistically significant difference between early (G1) and late (G5) trials (Fig. 2L). These findings suggest that reward approach-related population dynamics in pPVT^D2(−)^ neurons might not be sensitive to hunger states. To further assess our conclusions drawn from summary statistics, we again used FLMM to test the effects of latency and trial order on trial level responses (Supp. Fig. 3). In agreement with our earlier conclusions, FLMM revealed no association between either satiety nor vigor-related metrics and signal dynamics in pPVT^D2(−)^ neurons (Supp. Fig. 3C, D).

**Figure 2.**
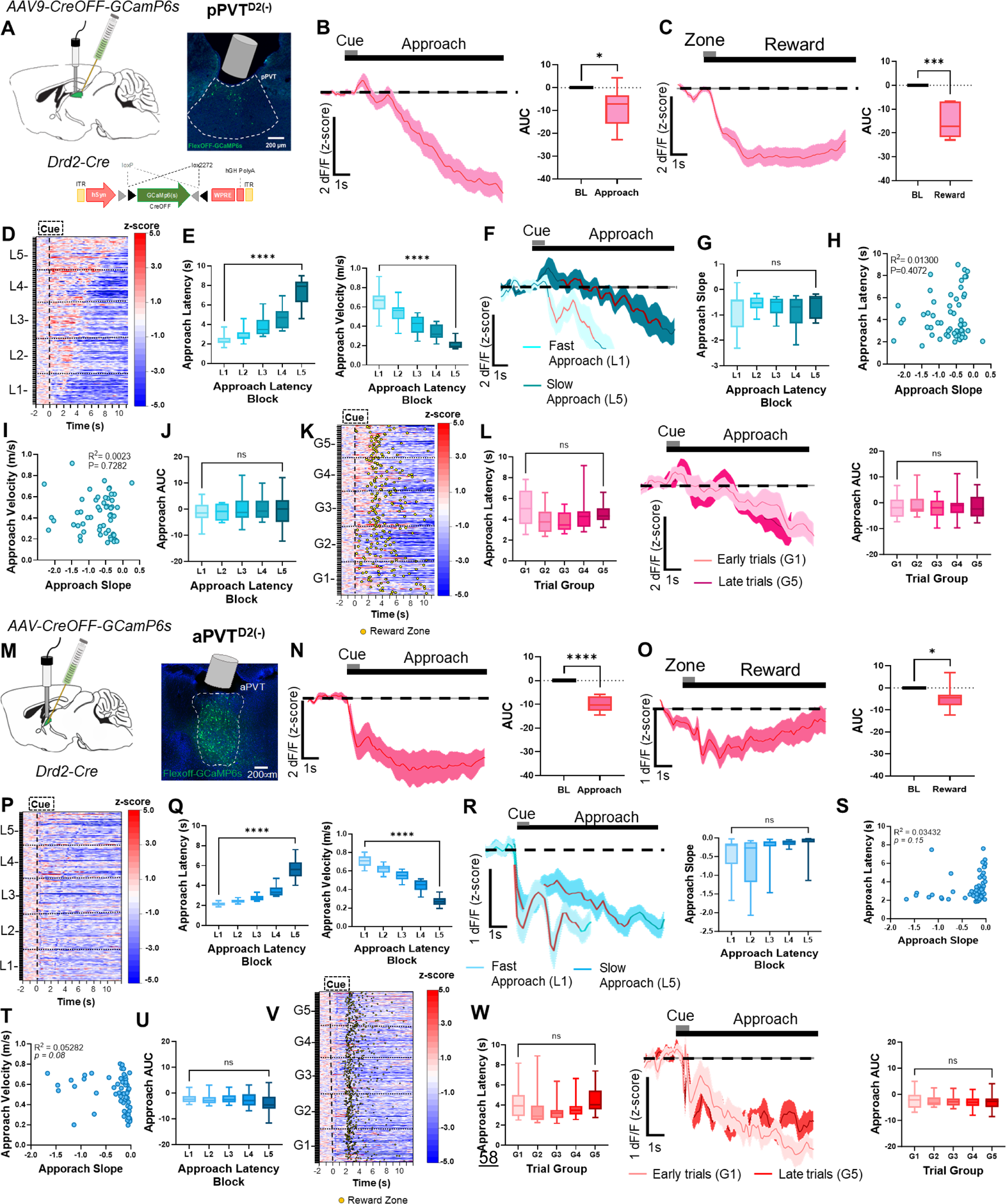
*In-vivo* activity dynamics of pPVT^D2(−)^ neurons. **(A)** *Left*: Schematic of the stereotaxic injections for selective expression of Cre-OFF GCaMP6s viral vectors and fiber implantation to target D2(−) neurons in the pPVT. *Right*: Representative image from a Drd2-Cre mouse expressing Cre-OFF GCaMP6s in the pPVT. **(B)** *Left*: Average GCaMP6s response from pPVT^D2(−)^ neurons during reward approach. *Right*: Quantification of the approach-evoked changes in GCaMP6s fluorescence in pPVT^D2(−)^ neurons. AUC, n = 420 trials from 6 mice, two-tailed paired t-test, \**p<0.05*. **(C)** *Left*: Average GCaMP6s responses from pPVT^D2(−)^ neurons during reward delivery. *Right*: AUC quantification of reward-evoked changes in GCaMP6s fluorescence in pPVT^D2(−)^ neurons. Two-tailed paired t-test, \*\**\*p<0.001*. **(D)** Heatmap showing approach responses from pPVT^D2(−)^ neurons, time-locked to cue onset, sorted by latency to approach the reward zone, and binned into 5 ‘approach latency blocks,’ L1, n = 79 trials, L2, n= 81 trials, L3, n = 84 trials, L4, n = 87 trials, L5, n = 89 trials from 6 mice. **(E)** *Left*: Latencies to reach the reward zone for each latency block. Repeated measures ANOVA *\*\*\*\*p<0.0001*. *Right*: Velocity (m/s) during reward approach calculated for each latency block. Repeated measures ANOVA, *\*\*\*\*p<0.0001*. **(F)** Average GCaMP6s dynamics for fast approach (L1) and slow approach (L5) in pPVT^D2(−)^ neurons. The red line indicates 20-80% of the slope of the line. **(G)** Slope-of-the-line quantifications of GCaMP6s activity from pPVT^D2(−)^ neurons across approach latency blocks. Repeated measures ANOVA, *p=0.25;* ns, not significant. **(H)** Correlation between the approach latency and the slope-of-the-line quantifications of GCaMP6s responses during reward approach. Each dot represents the GCaMP6s activity of individual mice averaged across approach latency blocks. **(I)** Correlation between the velocity (m/s) during the approach and the slope-of-the-line quantifications of GCaMP6s responses during the reward approach. **(J)** AUC quantification of the reward approach-evoked changes in GCaMP6s activity from pPVT^D2(−)^ neurons across approach latency blocks. Repeated measures ANOVA, *p=0.90;* ns, not significant. **(K)** Same as (D), but responses were sorted by trial order and binned into 5 ‘trial group blocks,’ G1, n = 79 trials, G2, n= 81 trials, G3, n = 84 trials, G4, n = 87 trials, G5, n = 89 trials from 6 mice. Yellow dots represent reward zone arrival. **(L)** *Left*: Latencies to reach the reward zone across trial group blocks. Repeated measures ANOVA, *p=0.11;* ns, not significant*. Middle*: Average GCaMP6s responses from pPVT^D2(−)^ neurons comparing approach trials performed early and late in the testing session. *Right*: AUC quantification of the reward approach-evoked changes in GCaMP6s activity from pPVT^D2(−)^ neurons across trial group blocks. Repeated measures ANOVA, *p=0.93*; ns, not significant. **(M)** *Left*: Schematic of stereotaxic injections and fiber implantation. *Right:* Representative image of D2(−) neurons in the aPVT. **(N)** *Left*: Average GCaMP6s response from aPVT^D2(−)^ neurons showing decreases in activity dynamics during reward approach. *Right:* AUC quantification of approach-evoked changes in GCaMP6s fluorescence in aPVT^D2(−)^ neurons. AUC, n = 650 trials from 8 mice, two-tailed paired t-test, *\*\*\*\*p<0.0001*. **(O)** *Left*: Average GCaMP6s response from aPVT^D2(−)^ neurons showing decreased activity dynamics during reward delivery*. Right:* AUC quantification of reward-evoked changes in GCaMP6s fluorescence in aPVT^D2(−)^ neurons. Two-tailed paired t-test, \**p<0.05*. **(P)** Heatmap showing approach responses of aPVT^D2(−)^ neurons, time-locked to cue onset, sorted by latency to approach the reward zone, and binned into 5 ‘approach latency blocks’ (L1 – L5), L1, n = 129 trials, L2, n= 129 trials, L3, n = 129 trials, L4, n = 129 trials, L5, n = 139 trials from 8 mice. **(Q)** *Left*: Latencies to reach the reward zone in seconds (s) for each approach latency block. Repeated measures ANOVA, *\*\*\*\*p<0.0001*. *Right:* Velocity (m/s) during reward approach calculated for each approach latency block. Repeated measures ANOVA, *\*\*\*\*p<0.0001*. **(R)** *Left*: Average GCaMP6s responses for fast and slow reward approach. The red line indicates 20-80% of the slope of the line. *Right:* Slope-of-the-line quantifications of GCaMP6s activity from aPVT^D2(−)^ neurons across approach latency blocks. Repeated measures ANOVA, *p=0.08*; ns, not significant **(S)** Correlation between approach latency and the slope-of-the-line quantifications per animal for each latency block. **(T)** Correlation between approach velocity and the slope-of-the-line of the line quantifications **(U)** AUC quantification during reward approach across latency blocks. Repeated measures ANOVA, \**p<0.05;* L1 vs. L5 Tukey’s multiple comparisons test, *p=0.14*. **(V)** Same as (P), but responses were sorted by trial order and binned into 5 ‘trial group blocks’ (G1 – G5). Yellow dots represent reward zone arrival. **(W)** *Left*: Quantification of latency of reward approach across trial group blocks. Repeated measures ANOVA, \**p<0.05;* G1 vs. G5 Tukey’s multiple comparisons test, *p=0.99*. *Middle:* Average GCaMP6s response for aPVT^D2(−)^ neurons during trials performed early (G1) and late (G5) during the session. *Right:* AUC quantification of GCaMP6s activity during approach across trial group blocks. Repeated measures ANOVA, *p=0.29*; ns, not significant. All data in the figure are shown as mean ±s.e.m.

Previous studies have also suggested that the anterior PVT (aPVT), where PVT^D2(−)^ neurons are most abundant^15,16^, modulates food-seeking and feeding behavior and is innervated by hypothalamic neurons that signal satiety^6,25,26^. Therefore, in a subset of mice, we recorded aPVT^D2(−)^ neurons while animals performed in our foraging-like reward task (Fig, 2M). Our findings showed that, like pPVT^D2(−)^ neurons, GCaMP fluorescence in aPVT^D2(−)^ neurons significantly decrease during reward approach and reward zone entry (Fig. 2N, O), suggesting that PVT^D2(−)^ neurons are functionally similar irrespective of their antero-posterior location in the PVT. Moreover, grouping trials based on either latency and trial order revealed no associations between these parameters and signal (Fig. 2P-W), a conclusion that was further supported by FLMM analysis (Supp. Fig. 3E, F). It is important to note that our independent assessment of the activity dynamics of pPVT^D2(−)^ and aPVT^D2(−)^ neurons assumes that these two belong to a largely homogenous subpopulation of PVT neurons. To directly test this prediction, we used FLMM and included recording location (i.e., aPVT or pPVT) as a covariate in our model. These analyses revealed no major effect of recording location cue and reward zone-related signal dynamics (Supp. Fig. 3G), thus supporting the notion that genetically-defined subpopulations of the PVT display similar functional properties regardless of their location within the antero-posterior axis of the PVT^16^.

### PVT^D2(+)^ and PVT^D2(−)^ neuronal activity dynamics during trial termination

Previous studies, including those from our group, have established that PVT neuronal activation is crucial for food-seeking behavior^11,13,22,27^. We now provide evidence that pPVT^D2(+)^ neurons are likely implicated in this process. However, other studies have suggested that PVT neuron activation might also promote the suppression of reward-seeking behavior^6,28,29^. This led us to hypothesize that PVT^D2(−)^ neurons, albeit being suppressed during the initiation of goal pursuits, signal the termination of such behaviors. To test this prediction, we independently examined the *in vivo* dynamics of pPVT^D2(+)^ and PVT^D2(−)^ neurons during ‘trial termination,’ defined as the moment when mice completed reward consumption and began to return to the trigger zone to initiate another trial (Fig. 3A) (See Methods). Our analysis revealed that trial termination resulted in a reduction in pPVT^D2(+)^ neuronal activity (Fig. 3B) and significant increases in both pPVT^D2(−)^ and aPVT^D2(−)^ neuronal activity (Fig. 3C, D), supporting our hypothesis that while pPVT^D2(+)^ neurons signal reward approach, PVT^D2(−)^ neurons signal the termination of food-seeking behavior. Interestingly, when trials were sorted and binned by latency to return to the trigger zone, neither pPVT^D2(+)^, nor pPVT^D2(−)^ neuronal activity was modulated by whether mice performed a ‘slow’ or ‘fast’ return (Fig. 3E, F). Surprisingly, for aPVT^D2(−)^, there was not a main effect of return latency block, but pairwise comparisons between the slopes of L1 and L5 revealed significant differences in aPVT^D2(−)^ (Fig. 3G). These findings were further supported by FLMM analysis which revealed a lack of effect of return latency on pPVT^D2(+)^ and pPVT^D2(−)^ neuronal signaling for both trial termination and trigger zone entry (Supp. Fig. 3H, J), and a main effect of return latency on aPVT^D2(−)^ neuronal signal at both termination and trigger zone entry (Supp. Fig. 3L). Lastly, when trials were sorted and binned by trial order, we found no statistically significant modulation by the hunger state of the animals for either neuronal type (Fig. 3H-J) and these findings were further supported by FLMM (Supp. Fig. 3I, K, M). Altogether, our findings indicate that PVT^D2(−)^ neuronal activity, but not pPVT^D2(+)^, signals the termination of goal pursuits, and that aPVT^D2(−)^, but not pPVT^D2(−)^ neuronal activity is influenced by motivation-related metrics such as latency to return to the trigger zone after trial termination.

**Figure 3.**
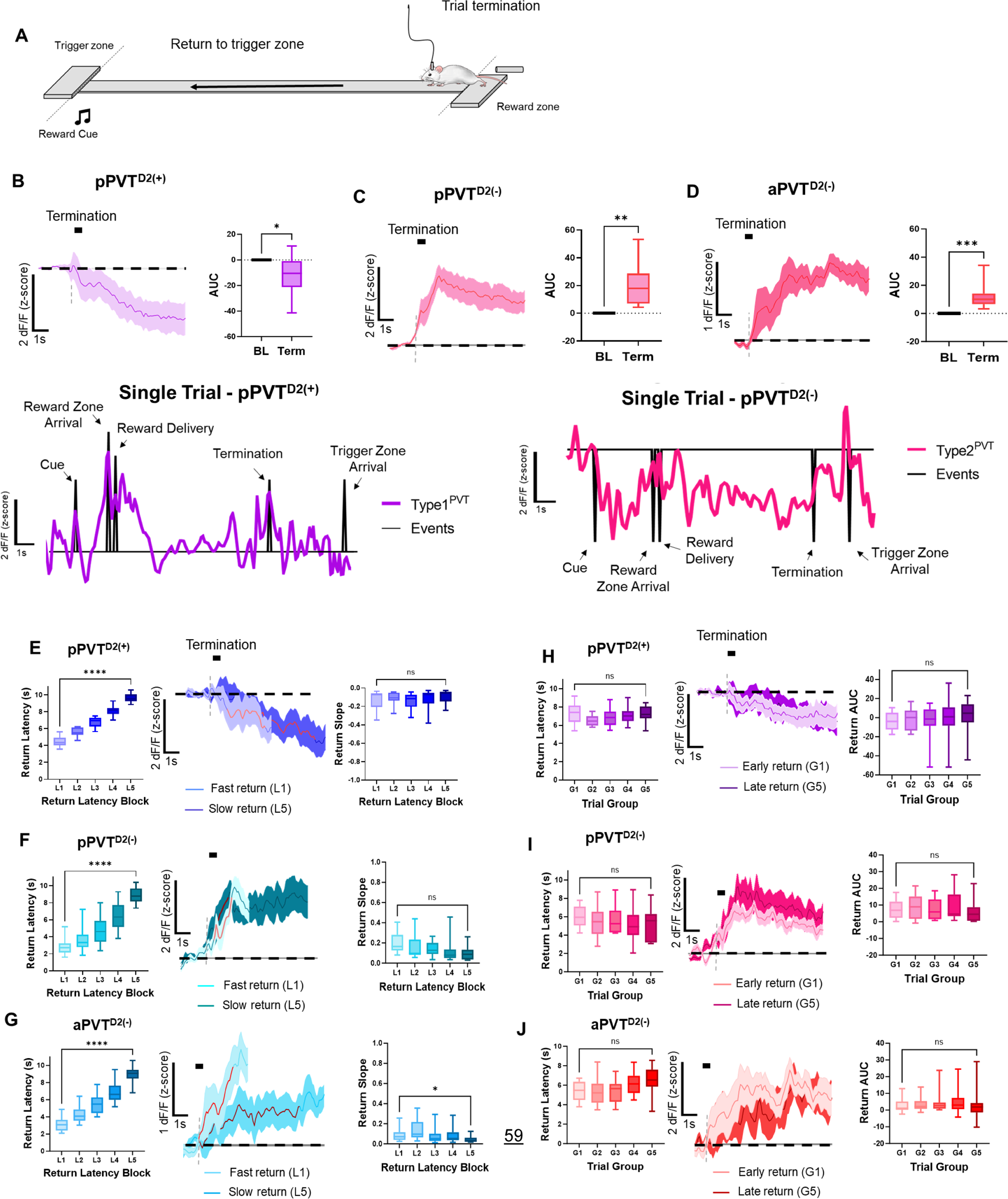
PVT^D2(+)^ and PVT^D2(−)^ neurons *in vivo* dynamics during trial termination. **(A)** Schematic depicting animal return to the trigger zone in our foraging-like reward-seeking task. **(B)** *Left*: Average GCaMP6s responses from pPVT^D2(+)^ neurons during trial termination and return. *Right:* Quantification of the return-evoked changes in GCaMP6s fluorescence in pPVT^D2(+)^ neurons. AUC, two-tailed paired t-test, \**p<0.05*. *Bottom:* In a complete trial, the activity dynamics of pPVT^D2(+)^ neurons from a sample subject during the reward foraging task are represented by the purple line, while the black ticks indicate various events within the trial. **(C)** *Left*: Average GCaMP6s responses from pPVT^D2(−)^ neurons during trial termination and return. *Right:* Quantification of the return-evoked changes in GCaMP6s fluorescence in pPVT^D2(−)^ neurons. AUC, two-tailed paired t-test, \*\**p<0.01*. *Bottom:* Same as (B) for pPVT^D2(−)^ neurons. **(D)** *Left*: Average GCaMP6s responses from aPVT^D2(−)^ neurons during trial termination and return. *Right:* Quantification of the return-evoked changes in GCaMP6s fluorescence in aPVT^D2(−)^ neurons. AUC, two-tailed paired t-test, \*\*\**p<0.001*. **(E)** *Left:* Latencies to reach the trigger zone across approach latency blocks for pPVT^D2(+)^ neuronal imaging. Repeated measures ANOVA, \*\*\*\**p<0.01. Middle:* Average GCaMP6s dynamics for fast return (L1) and slow return (L5) in pPVT^D2(+)^ neurons. The red line indicates 20-80% of the slope of the line. *Right:* Slope-of-the-line quantification of GCaMP6s return activity in pPVT^D2(+)^ neurons. Repeated measures ANOVA, *p=0.45*; ns, not significant. **(F)** *Left:* Latencies to reach the trigger zone across approach latency blocks for pPVT^D2(−)^ photometry recordings. Repeated measures ANOVA, \*\*\*\**p<0.01. Middle:* Average GCaMP6s dynamics for fast return (L1) and slow return (L5) in pPVT^D2(−)^ neurons. The red line indicates 20-80% of the slope of the line. *Right:* Slope-of-the-line quantification of GCaMP6s return activity in pPVT^D2(−)^ neurons across latency blocks. Repeated measures ANOVA, *p=0.28*; ns, not significant. **(G)** *Left:* Latencies to reach the trigger zone across approach latency blocks for aPVT^D2(−)^ photometry recordings. Repeated measures ANOVA, \*\*\*\**p<0.01. Middle:* Average GCaMP6s dynamics for fast return (L1) and slow return (L5) in aPVT^D2(−)^ neurons. The red line indicates 20-80% of the slope of the line. *Right:* Slope-of-the-line quantification of GCaMP6s return activity in aPVT^D2(−)^ neurons across latency blocks. Repeated measures ANOVA, *p=0.13;* G1 vs. G5 Tukey’s multiple comparisons test. **(H)** *Left:* Latencies to reach the trigger zone across trial group blocks for pPVT^D2(+)^ photometry recordings. Repeated measures ANOVA, *p=0.29;* ns, not significant. *Middle:* Average GCaMP6s return dynamics from pPVT^D2(+)^ neurons comparing return trials performed early and late in the testing session. *Right:* AUC quantification of GCaMP6s activity during the return in pPVT^D2(+)^ neurons across trial group blocks. Repeated measures ANOVA, *p=0.71*; ns, not significant. **(I)** *Left:* Latencies to reach the trigger zone across trial group blocks for pPVT^D2(−)^ photometry recordings. Repeated measures ANOVA, *p=0.18;* ns, not significant. *Middle:* Average GCaMP6s return dynamics in pPVT^D2(−)^ neurons for early and late trials. *Right:* AUC quantification of GCaMP6s activity during return in pPVT^D2(−)^ neurons across trial group blocks. Repeated measures ANOVA, *p=0.76*; ns, not significant. **(J)** *Left:* Latencies to reach the trigger zone across trial group blocks for aPVT^D2(−)^ photometry recordings. Repeated measures ANOVA, \**p<0.05;* G1 vs. G5 Tukey’s multiple comparisons test, *p=0.16*. *Middle:* Average GCaMP6s return dynamics in aPVT^D2(−)^ neurons for early and late trials. *Right:* AUC quantification of GCaMP6s activity during return in aPVT^D2(−)^ neurons across trial group blocks. Repeated measures ANOVA, *p=0.54*; ns, not significant. All data in the figure are shown as mean ±s.e.m.

### pPVT^D2(+)^–NAc and aPVT^D2(−)^–NAc terminals display similar activity dynamics to pPVT^D2(+)^ and PVT^D2(−)^ neurons

Approximately 80% of PVT neurons project to the NAc^30^, including both PVT^D2(+)^ and PVT^D2(−)^ neuronal subtypes^4,16,31^. Of note, the NAc plays a critical role in the learning and execution of goal-oriented behaviors^18,32–34^. For this reason, we next investigated how calcium signals recorded from the terminals of PVT^D2(+)^ and PVT^D2(−)^ neurons over the NAc correlate with task and behavior variables in our reward foraging-like behavioral task (Fig. 4). First, we used the same clustering analyses described earlier to compare the activity dynamics of these terminals between trials with different latencies and those at different stages of the session (Supp. Fig. 1H). Our results demonstrate that pPVT^D2(+)^–NAc terminals mirror the activity dynamics of pPVT^D2(+)^ neurons (Fig. 4A - U). Specifically, we observed robust increases in the fluorescence of pPVT^D2(+)^–NAc terminals during the reward approach, followed by a bi-phasic response at reward delivery (Fig. 4A - U). In addition, pPVT^D2(+)^–NAc terminals showed higher GCaMP signal ramps during fast trials compared to slow trials (Fig. 4D - U). We also found a negative correlation between approach latency and the slope of GCaMP signals, as well as a positive correlation between velocity to approach the reward and the slope of the GCaMP signal (Fig. 4G, H). This relationship between latency and signal was corroborated by FLMM (Supp. Fig. 3O). Surprisingly, and unlike for pPVT^D2(+)^ cell bodies, by assessing average signals across trial order groups we did not observe a statistically significant difference in the GCaMP signal of pPVT^D2(+)^–NAc terminals between early and late trials (Fig. 4J - U). However, FLMM analysis of pPVT^D2(+)^–NAc terminal recordings revealed a trial order effect at approximately 1s following reward zone entry (Supp. Fig. 3P). This discrepancy between our summary statistics analysis and FLMM, likely results from the capacity of the latter to assess significant associations at any given timepoint during each trial^23^. We also found that pPVT^D2(+)^–NAc terminals showed a significant decrease in activity during trial termination (Fig. 4L) and FLMM analysis revealed both latency and trial order effects after trial termination and trigger zone entry (Supp. Fig. 3S, T). Collectively, these results suggest that decreases in pPVT^D2(+)^–NAc terminal activity are associated with trial termination and/or task disengagement, and that termination-associated responses are modulated by both motivation and satiety.

**Figure 4.**
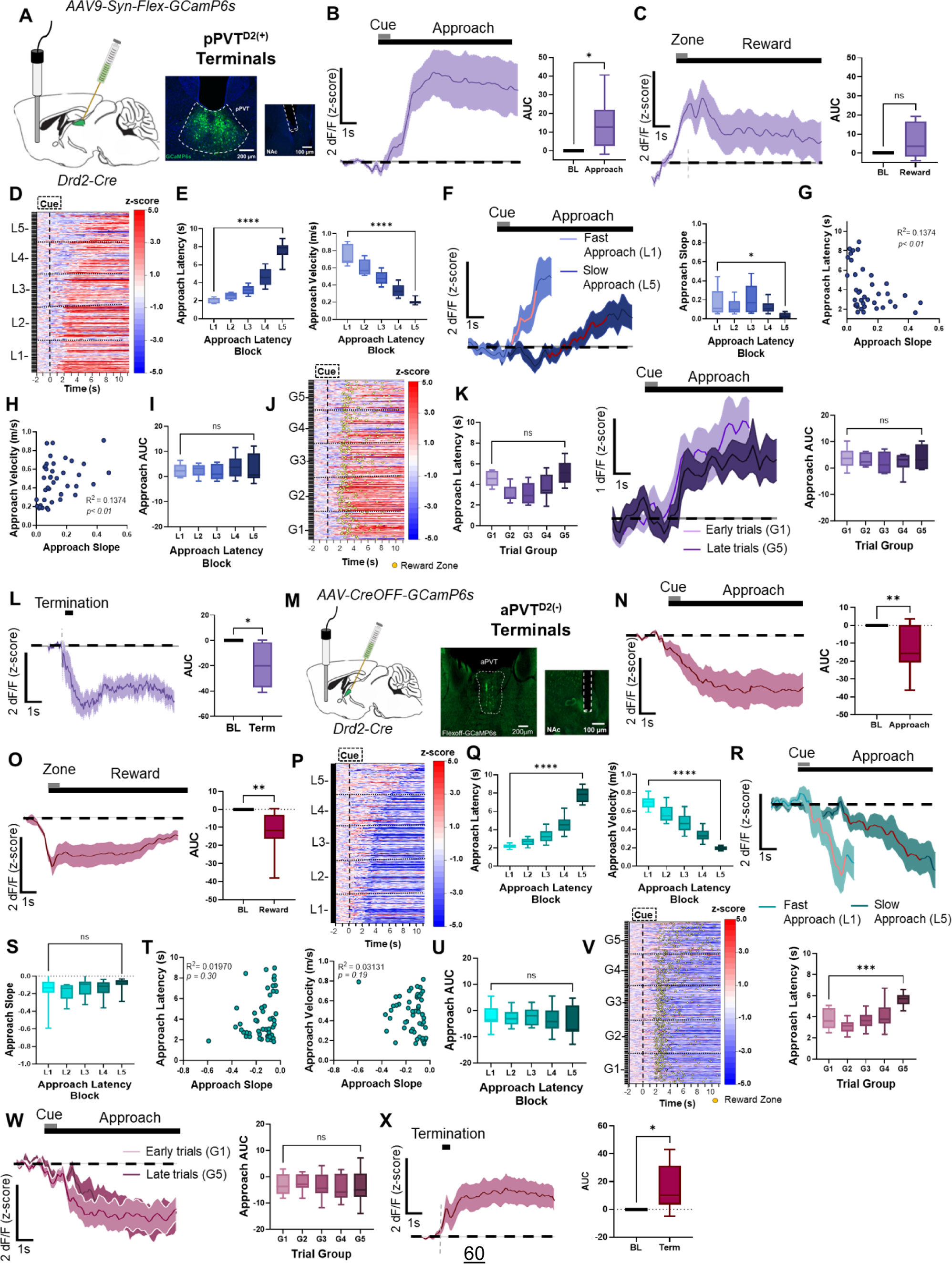
*In vivo* activity dynamics of PVT^D2(+)^–NAc and PVT^D2(−)^–NAc axon terminals. **(A)** *Left*: Schematic of stereotaxic injections and fiber implantation. *Right:* Representative images of GCaMP6s expression in D2(+) neurons in the pPVT and fiber placement in NAc. **(B)** *Left*: Average GCaMP6s approach-evoked responses from pPVT^D2(+)^ terminals during the reward approach. *Right*: Quantification of the approach-evoked changes in GCaMP6s fluorescence in pPVT^D2(+)^ terminals. AUC, n = 330 trials from 4 mice, two-tailed paired t-test, \**p<0.05*. **(C)** *Left*: GCaMP6s reward-evoked response in pPVT^D2(+)^ terminals during reward delivery. *Right*: Quantification of the reward-evoked changes in GCaMP6s fluorescence in pPVT^D2(+)^ terminals. AUC, two-tailed paired, *p=0.08*; ns, not significant. **(D)** Heatmap showing excitatory reward approach responses of pPVT^D2(+)^ terminals, time-locked to cue onset, sorted by latency to approach the reward zone, and binned into 5 ‘approach latency blocks’ (L1 – L5). L1, n = 65 trials, L2, n= 65 trials, L3, n = 65 trials, L4, n = 65 trials, L5, n = 70 trials from 4 mice. **(E)** *Left*: Latency to approach reward in seconds for pPVT^D2(+)^ terminals across latency blocks. Repeated measures ANOVA, *\*\*\*\*p<0.0001. Right*: Velocity (m/s) during reward approach calculated across each latency block. Repeated measures ANOVA, *\*\*\*\*p<0.0001*. **(F)** *Left*: Average GCaMP6s pPVT^D2(+)^ terminal dynamics for fast approach (L1) and slow approach (L5). The red line indicates 20-80% of the slope of the line. *Right*: Slope-of-the-line quantifications of GCaMP6s activity from PVT^D2(+)^ terminals across approach latency blocks. Repeated measures ANOVA, \**p<0.05*. **(G)** Correlation between the approach latency and the slope-of-the-line quantifications of GCaMP6s responses during reward approach in pPVT^D2(+)^ terminals. **(H)** Correlation between the approach velocity and the slope-of-the-line during the reward approach. **(I)** AUC quantification of the reward approach-evoked changes in GCaMP6s activity across approach latency blocks. Repeated measures ANOVA, *p=0.52;* ns, not significant. **(J)** Same as (D) but sorted by trial order and binned into 5 ‘trial group blocks’ (G1-G5). Yellow dots represent reward zone arrival. **(K)** *Left*: Latency to approach reward in seconds for pPVT^D2(+)^ terminals across latency blocks. Repeated measures ANOVA, *\*\*\*\*p<0.0001. Middle*: Average GCaMP6s approach dynamics of pPVT^D2(+)^ terminals comparing approach trials performed early and late in the session. *Right*: Max peak quantification of the approach-evoked changes in GCaMP6s activity from pPVT^D2(+)^ terminals across trial group blocks. Repeated measures ANOVA, *p=0.37*; ns, not significant. **(L)** *Left*: Average GCaMP6s responses from pPVT^D2(+)^ terminals during trial termination and return. *Right*: Quantification of the return-evoked changes in GCaMP6s fluorescence in pPVT^D2(+)^ terminals. AUC, two-tailed paired t-test, *\*p<0.05*. **(M)** *Left*: Schematic of stereotaxic injections and fiber implantation. *Right:* Representative images of GCaMP6s expression in D2(−) neurons in the aPVT and fiber placement in NAc. **(N)** *Left*: Average GCaMP6s approach-evoked responses from aPVT^D2(−)^ terminals during the reward approach. *Right*: Quantification of the approach-evoked changes in GCaMP6s in aPVT^D2(−)^ terminals. AUC, n = 646 trials from 6 mice, two-tailed paired t-test, *\*\*p<0.01*. **(O)** *Left*: Average GCaMP6s responses from aPVT^D2(−)^ terminals during reward delivery. *Right*: Quantification of reward-evoked changes in GCaMP6s. AUC, two-tailed paired t-test, *\*\*p<0.01*. **(P)** Heatmap showing inhibitory approach responses from aPVT^D2(−)^ terminals, time-locked to cue onset, sorted by latency to approach the reward zone, and binned into 5 ‘approach latency blocks’ (L1 – L5). L1, n = 127 trials, L2, n= 129 trials, L3, n = 127 trials, L4, n = 130 trials, L5, n = 133 trials from 6 mice. **(Q)** *Left*: Latencies to reach the reward zone in seconds for each approach latency block. Repeated measures ANOVA, *\*\*\*\*p = 0.0001*. *Right*: Velocity (m/s) during reward approach calculated for each approach latency block. Repeated measures ANOVA, *\*\*\*\*p<0.0001*. **(R)** Average GCaMP6s dynamics for aPVT^D2(−)^ terminals during fast and slow trials. **(S)** Slope-of-the-line quantifications of GCaMP6s activity from aPVT^D2(−)^ terminals across approach latency blocks. Repeated measures ANOVA, *p=0.42*; ns, not significant. **(T)** *Left*: No correlation between the approach latency and the slope-of-the-line quantifications of GCaMP6s responses from aPVT^D2(−)^ terminals during the approach. *Right*: No correlation between approach velocity and the slope-of-the-line quantifications of GCaMP6s responses during the reward approach. **(U)** AUC quantification of the approach-evoked changes in GCaMP6s activity across approach latency blocks. Repeated measures ANOVA, *p=0.11;* ns, not significant. ns, not significant. **(V)** *Left*: Same as (P), but responses were sorted by trial order and binned into 5 ‘trial group blocks’ (G1 – G5). Yellow dots represent reward zone arrival. *Right*: Latencies to reach reward across trial group blocks for aPVT^D2(−)^ terminals. Repeated measures ANOVA, *\*\*\*p<0.001*. **(W)** *Left*: Average GCaMP6s aPVT^D2(−)^ terminal dynamics comparing trials performed early and late in the session. *Right*: AUC quantification of the approach-evoked changes in GCaMP6s activity from aPVT^D2(−)^ terminals across trial group blocks. Repeated measures ANOVA, *p=0.26*; ns, not significant. **(X)** *Left*: Average GCaMP6s responses from aPVT^D2(−)^ terminals during trial termination and return. *Right*: Quantification of the return-evoked changes in GCaMP6s fluorescence in aPVT^D2(−)^ terminals. AUC, two-tailed paired t-test, *\*p<0.05*. All data in the figure are shown as mean ±s.e.m.

Finally, analysis of aPVT^D2(−)^–NAc terminal activity demonstrated similar dynamics to the broader aPVT^D2(−)^ neuronal population (Fig. 4M – X). Specifically, aPVT^D2(−)^–NAc terminals displayed significant decreases in activity during the reward approach and delivery (Fig. 4N, O). However, analysis of average signals for each trial type showed no significant differences in the activity of aPVT^D2(−)^–NAc terminals between fast vs. slow approach or early vs. late trials (Fig.4P-W). Lastly, trial termination resulted in increases in the activity of aPVT^D2(−)^–NAc terminals (Fig. 4X). To gain further insight into the association of parameters such as latency and trial order on the approach and termination dynamics of aPVT^D2(−)^–NAc terminal we used FLMM. Like for aPVT^D2(−)^ neurons, FLMM showed no effect of trial order on signal magnitude during reward approach (Supp. Fig. 3R). However, an effect of latency on signal was observed ahead of reward zone entry (Supp. Fig. 3Q). FLMM also revealed an association between latency to return to the trigger zone, but not trial order, and signal for aPVT^D2(−)^–NAc terminals (Supp. Fig. 3U, V), suggesting that like aPVT^D2(−)^ cell bodies, these terminals are sensitive to motivation-related variables. Collectively, our findings strongly suggest that motivation-related features and the encoding of goal-oriented actions of pPVT^D2(+)^ and PVT^D2(−)^ neurons are being relayed to the NAc through their respective terminals.

## DISCUSSION

We have identified two parallel thalamo-striatal pathways, namely PVT^D2(+)^–NAc and PVT^D2(−)^–NAc, whose *in vivo* dynamics respectively encode the initiation and termination of goal pursuits. In addition, we found that activity in these pathways, particularly the PVT^D2(+)^–NAc, mirrors that of the broader PVT^D2(+)^ and PVT^D2(−)^ neuronal populations and tracks the motivational state of subjects. These results are consistent with the notion that the PVT–NAc pathway guides motivated behaviors and offer a framework for interpreting seemingly contradictory findings from the literature, where manipulations of the PVT either increase or decrease reward seeking^6,10,11,13,16,22,25–29,35–38^. Indeed, our studies support the notion that the dorsal midline thalamus integrates signals about need states to shape goal-oriented behaviors^8,12,14,27,39^. Below, we discuss in detail our major findings and frame them within the broader context of the literature.

The PVT has rapidly emerged as an important node guiding motivated behaviors, a process that is thought to be mediated by its ability to integrate interoceptive signal and cognitive information to guide goal-oriented actions^8,12,14,27,39^. For instance, recent research has proposed that the activity of PVT neurons is modulated by hypothalamic-derived neuromodulators such as orexin and neuropeptide Y to promote need states and drive approach behaviors^13,40^. The PVT’s role in guiding both appetitive and aversive-motivated goal-oriented actions is thought to be largely dependent on its projections to the NAc^1,4,5,10,11,37,41^. Of note, some studies have suggested that state-dependent increases in PVT–NAc signaling are directly proportional to increased goal seeking behavior^1,5,11,13,22^. For example, studies investigating single-cell PVT–NAc activity have shown that putative inhibitory hypothalamic projections to the PVT are relaying licking information^10^. These inhibitory hypothalamic inputs could be sending information to the PVT regarding the hunger state of the animals. In line with these, NAc projecting PVT neurons have also been shown to develop cues responses which have been shown to depend on the animal’s hunger state and hypothalamic derived signals such as orexin^10,13,40^. These findings support the notion that need-based signals are conveyed to the PVT to regulate motivated behaviors in response to changes in internal states. On the other hand, others have argued that this relationship is rather inverse, for example, recent studies have shown that aPVT neurons^6,25^ and a subset of pPVT neurons^28,42^ mediate inappropriate reward-seeking behavior. Also, optogenetic activation of such PVT neurons inhibited active lever pressing in a sucrose-seeking task^6,28,29,42^. Here, we propose that thalamo-striatal projections arising from molecularly distinct subpopulations of the PVT are responsible for such seemingly contradictory observations. Specifically, we suggest that the striatal projections of PVT^D2(+)^ and PVT^D2(−)^ neurons, respectively, promote the initiation and termination of goal pursuits (but see ^42^). Our model appears in line with the differential anatomical distribution of PVT projections to the NAc^30^, and the known differential contribution of NAc subregions to goal-oriented behaviors^32,33,43,44^. In a recent publication, it was shown that while PVT^D2(+)^ neurons predominantly innervate the core and ventromedial shell of the NAc, the projections of PVT^D2(−)^ neurons largely concentrate in the dorsomedial shell and appear to be absent from the core^16^. Indeed, in the present study, we aimed to target the medial shell, a region where the two subtypes overlap in the NAc. However, future studies should aim to investigate whether the anatomical dissociation of PVT^D2(+)^ and PVT^D2(−)^ afferents to the NAc underlies their differential involvement in shaping instrumental actions.

It is worth noting that functional studies using patch-clamp electrophysiology have also identified differential connectivity between the PVT and specific neuronal subclasses of the NAc. Indeed, PVT axons have been shown to innervate NAc D1 medium spiny neurons (MSNs), D2 MSNs, cholinergic interneurons, and parvalbumin interneurons^1,28,42,45–48^, and some studies have argued that the PVT’s connectivity with specific NAc subclasses is selective and/or modulated in an experience and/or-dependent manner^1,28,42,47,48^. Future studies should investigate whether selective connectivity between the PVT and specific NAc subtypes maps onto either PVT^D2(+)^ or PVT^D2(−)^-derive input, and how this differential connectivity influences diverse aspects of motivated behaviors.

One critical aspect of our study is that, in addition to assessing photometry responses using standard approaches that rely on summary statistics, we have used a novel statistical framework, termed FLMM^23^, which enables hypothesis testing at every trial time point and focuses on trial-level signals (no averaging). Of note, while FLMM helped validate some of our conclusions drawn from summary statistics, it also yielded more nuanced information particularly regarding the magnitude and temporal distribution of effects. For example, FLMM uncovered an inverse relationship between trial order and signal magnitude at cue onset for pPVT^D2(+)^ neurons that was initially missed when averaged responses across groups were assessed. In addition, FLMM enabled statistical comparisons of responses drawn from PVT^D2(−)^ neurons within different PVT subregions, suggesting that responses in this population are rather similar and largely independent of anatomical location. Thus, our findings support the notion that FLMM is a robust statistical framework that facilitates extracting meaningful observations from fiber photometry signals^23^.

Here, we have shown that pPVT^D2(+)^ neurons and their projections to the NAc display similar response profiles and mirror motivational variables such as behavioral vigor and sensitivity to satiety levels. This observation is consistent with prior studies showing that PVT neuronal responses to motivational cues are shaped by state dependent processes, as highlighted earlier^6,13^. It is worth noting that while the responses of PVT^D2(−)^ neurons were similarly consistent across anatomical location (aPVT vs pPVT) and striatal projections (aPVT^D2(−)^–NAc), some key differences were observed. Specifically, unlike pPVT^D2(−)^ neurons, both aPVT^D2(−)^ neurons and their projections to the NAc appeared to be sensitive to motivation-related metrics such as latency to initiate and terminate a goal-oriented action and level of satiety. Although it is possible that these differences reflect a unique capacity of the aPVT^D2(−)^ population to track state-dependent information, an alternative interpretation is that these discrepancies arise from the differences in the sample sizes included in our analyses. Indeed, our pPVT^D2(−)^ dataset includes one third fewer trials than both our aPVT^D2(−)^ and aPVT^D2(−)^–NAc datasets. Future studies should investigate whether these potential differences across PVT^D2(−)^ populations are biologically relevant.

It is intriguing that while serving complementary functions in the initiation and termination of goal-oriented behaviors, the activity of PVT^D2(+)^ and PVT^D2(−)^ neurons appears to be largely mutually exclusive, with PVT^D2(+)^ neurons activated at initiation when PVT^D2(−)^ activity is suppressed and PVT^D2(−)^ neurons becoming activated at termination when PVT^D2(+)^ activity is reduced. Similar opposing dynamics were observed in PVT subpopulations while mice engaged in mutually exclusive defensive responses^5^. While this pattern of activity suggests that an inhibitory relationship exists between these two neuronal classes, the rodent thalamus (including PVT) is largely devoid of interneurons^49,50^, and PVT^D2(+)^ and PVT^D2(−)^ neurons are not synaptically connected to one another^16^. One potential mechanism for mediating the inhibitory interaction between these two functional subclasses of the PVT is the thalamic reticular nucleus (TRN). The TRN is a thin sheet of GABAergic neurons that surrounds the thalamus and is known to play a key role in gating sensory information by modulating thalamo-cortical transmission^50,51^. Recent work confirms earlier anatomical models suggesting that the TRN mediates interactions between different thalamic nuclei^51–54^. Although largely speculative at present, in the context of the PVT, it is possible that the TRN plays a similar role in mediating interactions between the PVT^D2(+)^ and PVT^D2(−)^ pathways. Specifically, the TRN may act as a hub that receives input from both pathways and modulates their respective activities through inhibitory interactions, resembling an open-loop configuration^55,56^. Of note, recent experimental evidence supports the notion that an open-loop configuration involving PVT and TRN likely exists^57^. Such architecture could allow for fine-tuning of goal-oriented behavior (i.e., engagement and disengagement of actions) by ensuring that the appropriate PVT–NAc pathway is activated or inhibited at any given time. Future work should be aimed at empirically testing this prediction.

Although here we have shown that two major subtypes of PVT neurons, namely PVT^D2(+)^ and PVT^D2(−)^, differentially signal the initiation and termination of reward-seeking, likely through their projections to the NAc, our classification might have limited our ability to extract more nuanced information of how PVT subpopulations contribute to goal-oriented behaviors. Indeed, recent research using single-cell RNA sequencing has identified at least five potential molecularly distinct PVT subtypes^15^ (See also^19^). Importantly, the study concluded that a major node of separation across transcriptomically distinct PVT neurons lies in the expression of dopamine D2 receptor gene *Drd2*, such that PVT^D2(+)^ neurons can be further subdivided into two subtypes: *Esr1*^+^ and *Col12a1*^+^, while PVT^D2(−)^ can be separated into two additional subtypes: *Npffr1*^+^ and *Hcrtr1*^+^. A fifth subpopulation of PVT neurons is distributed across the antero-posterior axis of the PVT and expresses *Drd3*^15^. Of note, according to Gao et al., 2023, the five major PVT subclasses also display marked heterogeneity. Thus, while our findings provide valuable insights into the role of PVT subtypes in motivated behaviors, future studies should investigate the contributions of other PVT subpopulations to fully understand the complex dynamics of the PVT–NAc circuitry. For instance, it is possible that some of the functional differences in PVT^D2(−)^ neurons observed across the antero-posterior axis of the PVT map onto discrete subpopulations of *Drd2*-negative neurons.

Some additional limitations of our study deserve to be acknowledged. Here, we have monitored the activity of PVT^D2(+)^ and PVT^D2(−)^ neurons and how they relate to the performance of motivated behavior in mice. As such, we cannot conclude that the activity dynamics we have uncovered are causally related to the different aspects of goal-oriented behavior that they appear to correlate with. Moreover, due to the nature of the fiber photometry technique, it is possible that the observed responses in PVT neuronal activity we report here represent the combined responses of additional PVT subpopulations beyond PVT^D2(+)^ and PVT^D2(−)^ neurons. Indeed, while five molecularly distinct PVT cells were recently described, top markers within these subpopulations displayed minimal overlap with one another suggesting that additional cell-type heterogeneity likely exists within the PVT. Another important limitation is that our experiments were conducted exclusively during the light (inactive) phase. Considering that previous studies have demonstrated diurnal variability in the intrinsic activity and membrane properties of PVT neurons^58^, it would be important to investigate how time of day impacts the activity dynamics of PVT^D2(+)^ and PVT^D2(−)^ neurons described in our study. Finally, it is crucial to acknowledge that our study has demonstrated consistent responses among PVT^D2(−)^ neurons in our task, regardless of their antero-posterior location. However, it is important to note that our focus was exclusively on tracking the activity of PVT^D2(+)^ neurons in the pPVT. This decision stemmed from the sparse distribution of PVT^D2(+)^ neurons in the most anterior segments of the aPVT, as documented by Gao et al. (2020). While this approach was chosen for practical reasons, it is essential to highlight the absence of comparisons across the various antero-posterior locations of the PVT for this neuronal population.

The findings presented here shed light on the neural circuits that underlie motivated behaviors, including initiation, vigor, and termination. These processes are central to achieving goals and maintaining appropriate levels of motivation in everyday life. However, deficits in motivation are associated with a range of psychiatric conditions, such as substance abuse, binge eating, and anhedonia in depression^59,60^. Therefore, a deeper understanding of the neural basis of motivated behavior may offer new insights into the treatment of these debilitating conditions. By revealing the specific neuronal pathways involved in motivation and how they interact, this research may facilitate the identification of new therapeutic targets for interventions aimed at restoring healthy motivational processes in individuals with such conditions.

## ACKNOWLEDGEMENTS

We thank Dr. Kirstie Cummings, Dr. Jeremy Day, Dr. Ryan Lingg and Briana Machen for their comments on the manuscript. We also thank the NIHM Section on Instrumentation for their contribution to building the foraging maze apparatus, the NIMH Rodent Behavior Core for their assistance in the behavioral experiments, and the NIMH Systems Neuroscience Imaging Resource. This work was supported by K99/R00 MH126429 to S.B., a NARSAD Young Investigator Award by the Brain and Behavior Research Foundation to S.B., and by the NIMH Intramural Research Program (1ZIAMH002950) to M.A.P.

## AUTHOR CONTRIBUTIONS

S.B. performed and analyzed all experiments. I.K., A.B., and S.R.G. assisted with behavioral experiments. C.G. assisted with surgical techniques. G.L. and F.P. assisted with the FLMM analyses. I.K. and E. M. assisted with data analysis. I.K. assisted with histology procedures. S.B. and M.A.P. designed the study and interpreted the results. S.B. and M.A.P. wrote the paper.

## FIGURE LEGENDS

**Supplemental Figure 1.**
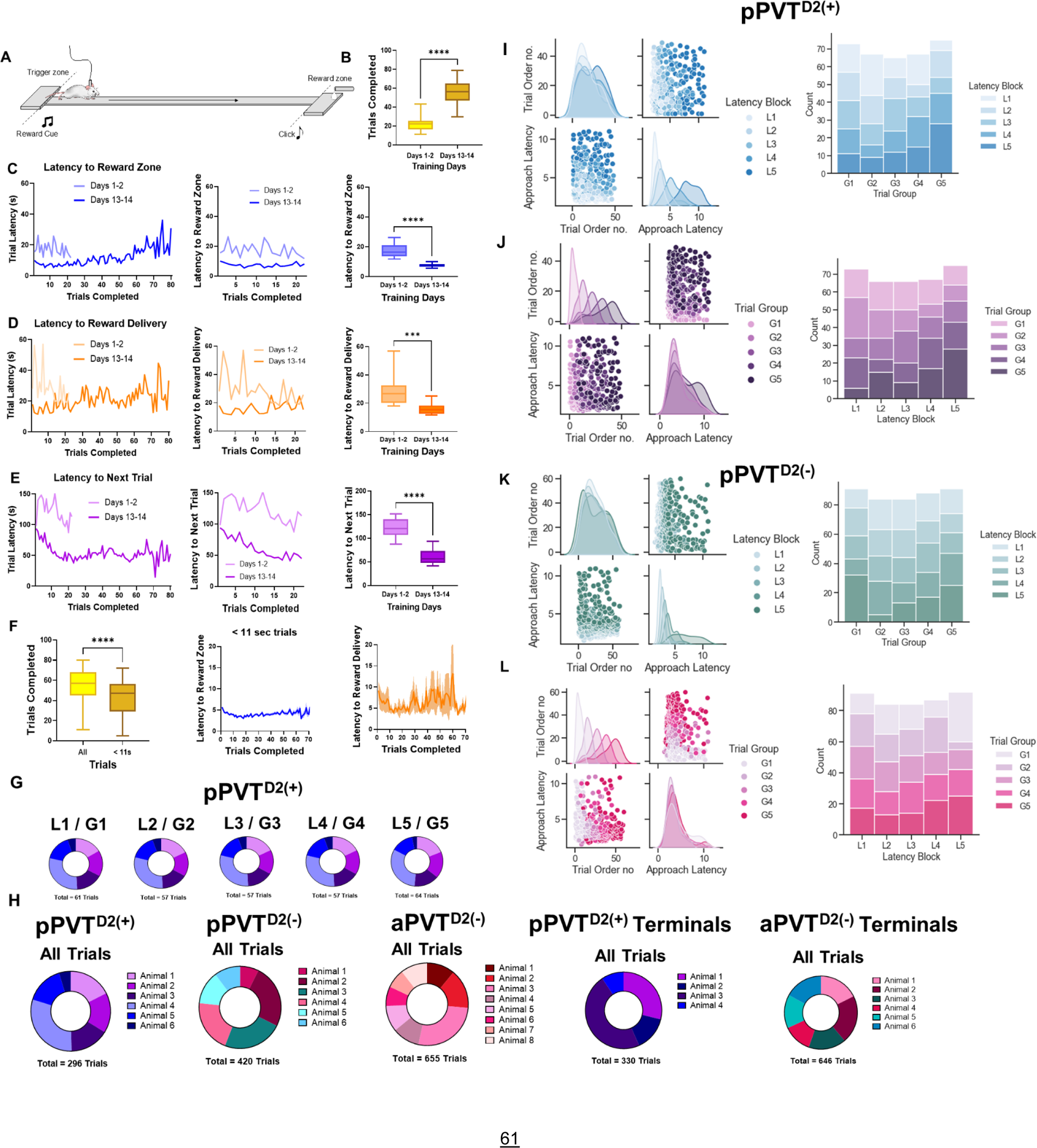
Mice performance in foraging–like task to characterize motivated behavior in rodents and trial distribution per experimental group. **(A).** Schematics depicting our foraging-like reward-seeking task. **(B)** Quantification of trials completed during the first days of training (Days 1-2) and the last days of training (Days 13-14) for all the mice included in the study. Two-tailed paired t-test, *\*\*\*\*p<0.0001*. **(C)** *Left:* Comparisons of the latencies to the reward zone. Lighter colors indicate the first days of training, and darker colors indicate the last days of training. *Middle:* Comparison of the latencies to reward zone between equivalent trials for training days 1-2 and days 13-14 (approx. 20 first trials). Right: Quantifications of (*Middle*). Two-tailed paired t-test, to zone \*\*\*\**p*<0.0001. **(D)** Same as (C) but for latencies to reward delivery. Two-tailed paired t-test, to delivery \*\*\**p*<0.001. **(E)** Same as (D) but for latencies to next trial. Two-tailed paired t-test, to next trial \*\*\*\**p*<0.0001. **(F)** *Left:* Comparison of all trials completed during testing with those included in the photometry analysis. On average, approximately ten trials were excluded per testing session. Two-tailed paired t-test, *\*\*\*\*p<0.0001*. *Middle:* Plots showing average latencies to the reward zone throughout the testing session for trials included in the photometry analysis. *Right:* Same as (Middle) but for latencies to reward delivery. **(G)** Pie charts showing the trial distribution per both approach latency blocks (L1-L5) and trial group blocks (G1-G5). **(H)** Pie chart showing individual trial contribution per animal for all groups tested in the study (pPVT^D2(+)^, pPVT^D2(−)^, aPVT^D2(−)^, pPVT^D2(+)^ -NAc terminals, aPVT^D2(−)^ -NAc terminals). **(I)** *Left:* Density estimates plots for trials during pPVT^D2(+)^ neuronal imaging and sorted by approach latency blocks. *Right*: Trial distribution for those trials performed during pPVT^D2(+)^ neuronal imaging showing proportion of trials in approach latency blocks and their distribution across trial group blocks. **(J)** *Left:* Density estimates graphs for trials during pPVT^D2(+)^ neuronal imaging and sorted by trial group blocks. *Right*: Trial distribution for those trials performed during pPVT^D2(+)^ neuronal imaging showing proportion of trials in trial group blocks and their distribution across approach latency blocks. **(K)** Same as (I) but for trials during pPVT^D2(−)^ neuronal imaging. **(L)** Same as (J) but for trials during pPVT^D2(−)^ neuronal imaging. All data in the figure are shown as mean ±s.e.m.

**Supplementary Figure 2.**
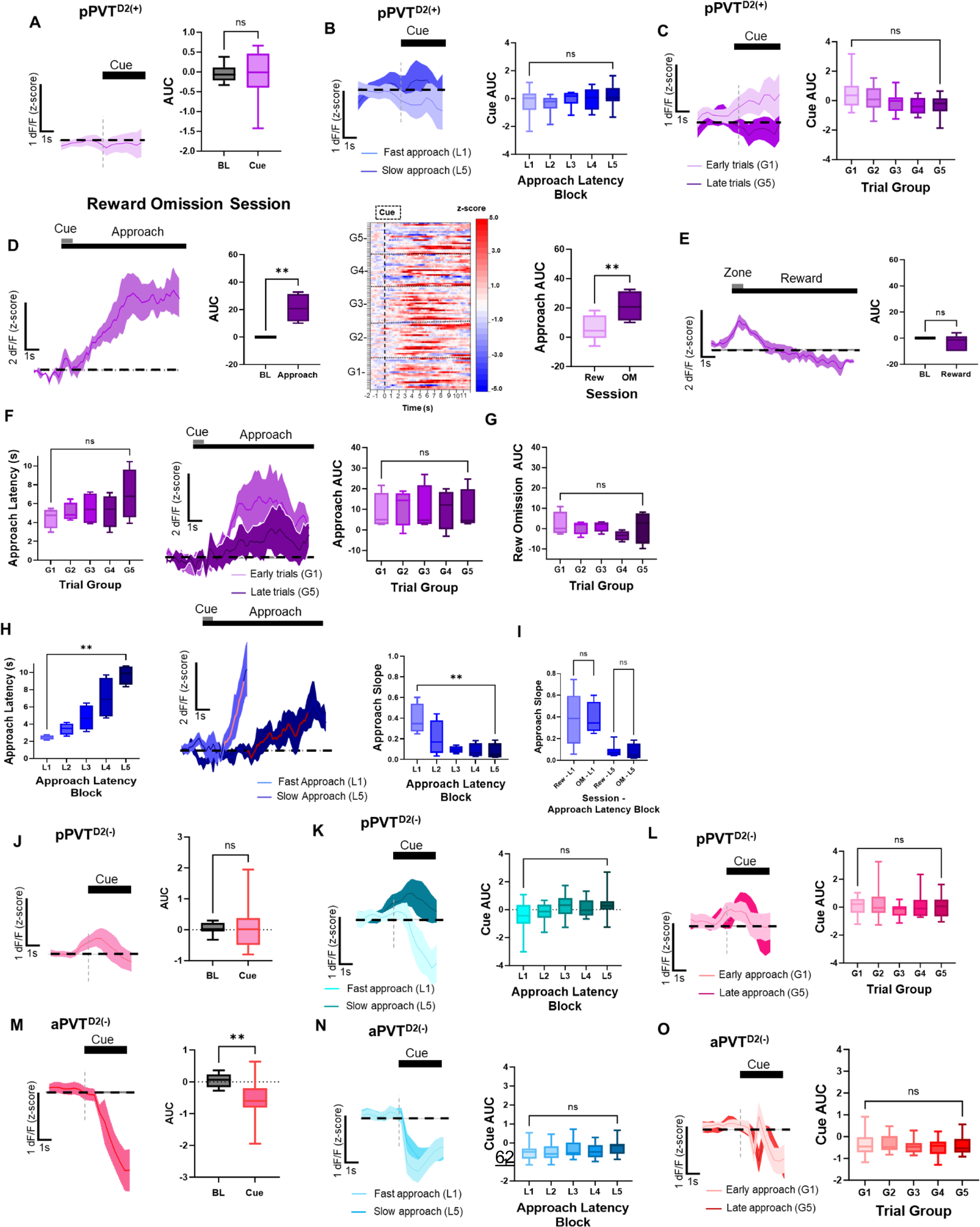
*In vivo* dynamics of PVT^D2(+)^ and PVT^D2(−)^ neurons during cue presentation and during reward omission testing session. **(A)** *Left:* Average approach-evoked GCaMP6s responses of pPVT^D2(+)^ neurons during cue presentation. *Right:* AUC quantification of baseline and cue activity of pPVT^D2(+)^ neurons. Two-tailed paired t-test, *p=0.82*; ns, not significant. **(B)** *Left:* Average GCaMP6s responses of pPVT^D2(+)^ neurons during cue presentation grouped by approach latency blocks. *Right:* AUC quantifications of GCaMP6s activity from pPVT^D2(+)^ neurons across approach latency blocks. Repeated measures ANOVA, *p=0.19;* ns, not significant. **(C)** Same as (B) but grouped by trial group blocks. Repeated measures ANOVA, \**p<0.05;* G1 vs. G5 Tukey’s multiple comparisons test, *p=0.13*. **(D)** *Left:* Average approach-evoked GCaMP6s responses of pPVT^D2(+)^ neurons in the OM session. *Middle Left:* AUC quantification of baseline and approach activity of pPVT^D2(+)^ neurons in the OM session. Two-tailed paired t-test, *\*\*p<0.01*. *Middle Right:* Heatmap of GCaMP6s responses from pPVT^D2(+)^ neurons during approach in the OM session. GCaMP6s responses were time-locked to cue onset, and trials were sorted by trial order and binned into 5 ‘trial group blocks’ (G1 – G5). *Right:* AUC quantification comparing approach-evoked GCaMP6s responses of pPVT^D2(+)^ neuronal responses between rewarded (Rew) and unrewarded (OM) testing sessions. Two-tailed unpaired t-test *\*\*p<0.01*. **(E)** *Left:* Average pPVT^D2(+)^ neuronal GCaMP6s responses when mice entered the food port but were not rewarded. *Right:* AUC quantification of the reward omission-evoked changes in GCaMP6s fluorescence in pPVT^D2(+)^ neurons. Two-tailed paired t-test, *p=0.24*; ns, not significant. **(F)** *Left:* Latencies to reach the reward zone across trial group blocks. Repeated measures ANOVA, *p=0.26;* ns, not significant*. Middle:* Average pPVT^D2(+)^ GCaMP6s responses during approach in the OM session for early and late trials. *Right:* AUC quantification of pPVT^D2(+)^ GCaMP6s activity in the OM session for trial group blocks. Repeated measures ANOVA, *p =0.71*; ns, not significant. **(G)** Same as (F) but for pPVT^D2(+)^ during reward omission. Repeated measures ANOVA, *p=0.44*; ns, not significant. **(H)** *Left:* Latencies to reach the reward zone in seconds for each approach latency block during the reward omission session. Repeated measures ANOVA, *\*\*p<0.0001. Middle:* Average pPVT^D2(+)^ neuronal GCaMP6s responses for fast and slow reward approach in the OM session. The red line indicates 20-80% of the slope of the line. *Right:* In the OM session, slope-of-the-line quantifications of pPVT^D2(+)^ neuronal GCaMP6s activity across approach latency blocks. Repeated measures ANOVA, *\*\*p<0.01*. **(I)** Slope-of-the-line quantifications of pPVT^D2(+)^ neuronal GCaMP6s activity comparing latencies during fast approach (L1) and slow approach (L5) between rewarded (Rew) and unrewarded (OM) testing sessions. L1-two-tailed unpaired t-test, *p=0.97*; ns, not significant. L5-two-tailed unpaired t-test, *p= 0.65*; ns, not significant. **(J)** *Left:* Average approach-evoked GCaMP6s responses of pPVT^D2(−)^ neurons during cue presentation. *Right:* AUC quantification of baseline and cue activity of pPVT^D2(−)^ neurons. Two-tailed paired t-test, *p=0.67*; ns, not significant. **(K)** *Left:* Average GCaMP6s responses of pPVT^D2(−)^ neurons during cue presentation grouped by approach latency blocks. *Right:* AUC quantifications of GCaMP6s activity from pPVT^D2(−)^ neurons across approach latency blocks. Repeated measures ANOVA, \**p<0.05;* L1 vs. L5 Tukey’s multiple comparisons test, *p=0.06*. **(L)** Same as (K) but grouped by trial group blocks. Repeated measures ANOVA, *p=0.50*; ns, not significant. **(M)** *Left:* Average approach-evoked GCaMP6s responses of aPVT^D2(−)^ neurons during cue presentation. *Right:* AUC quantification of baseline and cue activity of aPVT^D2(−)^ neurons. Two-tailed paired t-test, \*\**p<0.01*. **(N)** *Left:* Average GCaMP6s responses of aPVT^D2(−)^ neurons during cue presentation grouped by approach latency blocks. *Right:* AUC quantifications of GCaMP6s activity from aPVT^D2(−)^ neurons across approach latency blocks. Repeated measures ANOVA, *p=0.19*; ns, not significant. **(O)** Same as (N) but grouped by trial group blocks. Repeated measures ANOVA, *p=0.53*; ns, not significant. All data in the figure are shown as mean ±s.e.m.

**Supplementary Figure 3.**
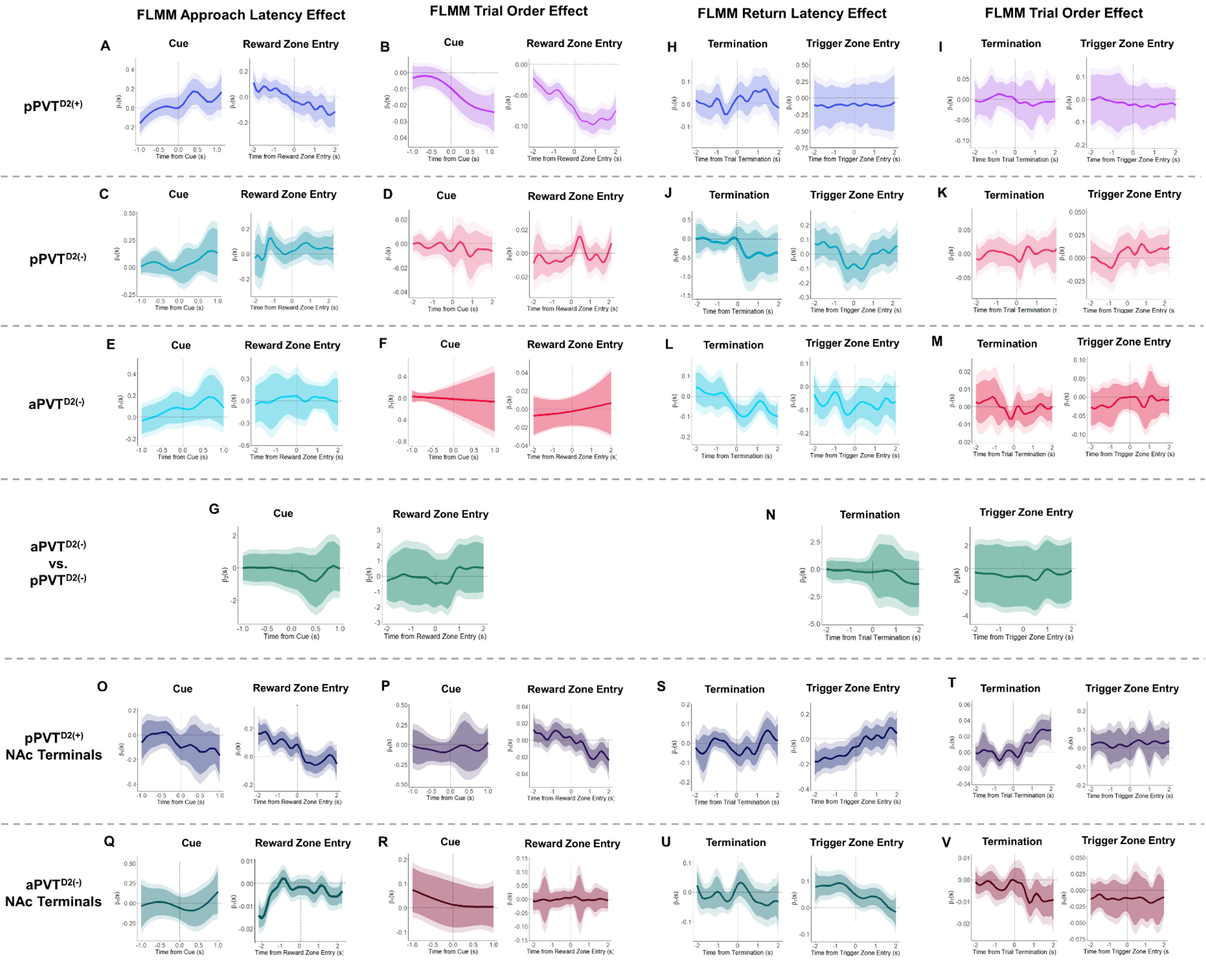
*In vivo* dynamics of PVT^D2(+)^ and PVT^D2(−)^ neurons and terminals using the novel FLMM analysis during distinct trial events. **(A)** FLMM coefficient estimates plots of the approach latency effect and statistical significance at each trial time-point results for the photometric responses of pPVT^D2(+)^ neurons for cue presentation and reward zone entry. No association between pPVT^D2(+)^ GCaMP6s responses and approach latency at cue presentation (*left*) nor at reward zone entry (*right*). **(B)** FLMM coefficient estimates plots of the approach trial order effect and statistical significance at each trial time-point results for the photometric responses of pPVT^D2(+)^ neurons for cue presentation and reward zone entry. The plots show a negative association between pPVT^D2(+)^ GCaMP6s responses and trial order right after cue presentation (*left*) and before reward zone entry (*right*). **(C)** Same as (A) but for photometric responses of pPVT^D2(−)^ neuron. No association between pPVT^D2(−)^ GCaMP6s responses and approach latency at cue presentation (*left*) or at reward zone entry (*right*). **(D)** Same as (B) but for photometric responses of pPVT^D2(−)^ neuron. No association between pPVT^D2(−)^ GCaMP6s responses and trial order at cue presentation (*left*) nor at reward zone entry (*right*). **(E)** Same as (A) but for photometric responses of aPVT^D2(−)^ neurons. No association between aPVT^D2(−)^ GCaMP6s responses and approach latency at cue presentation (*left*) or at reward zone entry (*right*). **(F)** Same as (B) but for photometric responses of aPVT^D2(−)^ neurons. No association between aPVT^D2(−)^ GCaMP6s responses and trial order at cue presentation (*left*) nor at reward zone entry (*right*). **(G)** FLMM coefficient estimates plots applying ‘recording location’ (i.e., aPVT or pPVT) as a covariate and showing statistical significance at each trial time-point results for the photometric responses of PVT^D2(−)^ neurons for cue presentation and reward zone entry. No statistically significant differences between aPVT^D2(−)^ GCaMP6s responses and pPVT^D2(−)^ GCaMP6s responses at cue presentation (*left*) nor at reward zone entry (*right*). **(H)** FLMM coefficient estimates plots of the return latency effect and statistical significance at each trial time-point results for the photometric responses of pPVT^D2(+)^ neurons for trial termination and trigger zone entry. No association between pPVT^D2(+)^ GCaMP6s responses and return latency at trial termination (*left*) nor at trigger zone entry (*right*). **(I)** FLMM coefficient estimates plots of the return trial order effect and statistical significance at each trial time-point results for the photometric responses of pPVT^D2(+)^ neurons for trial termination and trigger zone entry. No association between pPVT^D2(+)^ GCaMP6s responses and trial order at trial termination (*left*) nor at trigger zone entry (*right*). **(J)** Same as (H) but for photometric responses of pPVT^D2(−)^ neuron. No association between pPVT^D2(−)^ GCaMP6s responses and return latency at trial termination (*left*) nor at trigger zone entry (*right*). **(K)** Same as (I) but for photometric responses of pPVT^D2(−)^ neuron. No association between pPVT^D2(−)^ GCaMP6s responses and trial order at trial termination (*left*) nor at trigger zone entry (*right*). **(L)** Same as (H) but for photometric responses of aPVT^D2(−)^ neuron. No association between aPVT^D2(−)^ GCaMP6s responses and return latency at trial termination (*left*) nor at trigger zone entry (*right*). **(M)** Same as (I) but for photometric responses of aPVT^D2(−)^ neuron. No association between aPVT^D2(−)^ GCaMP6s responses and trial order at trial termination (*left*) nor at trigger zone entry (*right*). **(N)** FLMM coefficient estimates plots applying ‘recording location’ (i.e., aPVT or pPVT) as a covariate and showing statistical significance at each trial time-point results for the photometric responses of PVT^D2(−)^ neurons for trial termination and trigger zone entry. No statistically significant differences between aPVT^D2(−)^ GCaMP6s responses and pPVT^D2(−)^ GCaMP6s responses at trial termination (*left*) nor at trigger zone entry (*right*). **(O)** FLMM coefficient estimates plots of the approach latency effect and statistical significance at each trial time-point results for the photometric responses of pPVT^D2(+)^ terminals for cue presentation and reward zone entry. *Left:* No association between pPVT^D2(+)^ terminal responses and approach latency at cue presentation. *Right:* Plot showing a negative association between pPVT^D2(+)^ terminal responses and approach latency before reward zone entry. **(P)** FLMM coefficient estimates plots of the approach trial order effect and statistical significance at each trial time-point results for the photometric responses of pPVT^D2(+)^ terminals for cue presentation and reward zone entry. *Left:* No association between pPVT^D2(+)^ terminal responses and trial order at cue presentation. *Right:* Plot showing a negative association between pPVT^D2(+)^ terminal responses and trial order approximately 1 sec after reward zone entry. **(Q)** Same as (O) but for photometric responses of aPVT^D2(−)^ terminals. *Left:* No association between aPVT^D2(−)^ terminal responses and approach latency at cue presentation. *Right:* Plot showing a positive association between aPVT^D2(−)^ terminal responses and approach latency before reward zone entry. **(R)** Same as (P) but for photometric responses of aPVT^D2(−)^ terminals. No association between aPVT^D2(−)^ terminal responses and trial order at cue presentation (*left*) nor at reward zone entry (*right*). **(S)** FLMM coefficient estimates plots of the return latency effect and statistical significance at each trial time-point results for the photometric responses of pPVT^D2(+)^ terminals for trial termination and trigger zone entry. *Left:* No association between pPVT^D2(+)^ terminal responses and return latency at trial termination. *Right:* Plot showing a positive association between pPVT^D2(+)^ terminal responses and return latency before trigger zone entry. **(T)** FLMM coefficient estimates plots of the return trial order effect and statistical significance at each trial time-point results for the photometric responses of pPVT^D2(+)^ terminals for trial termination and trigger zone entry. *Left:* Plot showing a positive association between pPVT^D2(+)^ terminal responses and return trial order approx. 1 sec after trial termination. *Right:* No association between pPVT^D2(+)^ terminal responses and return trial order during trigger zone arrival. **(U)** Same as (S) but for photometric responses of aPVT^D2(−)^ terminals. *Left:* No association between aPVT^D2(−)^ terminal responses and return latency at trial termination. *Right:* Plot showing a negative association between aPVT^D2(−)^ terminal responses and return latency before trigger zone entry. **(V)** Same as (T) but for photometric responses of aPVT^D2(−)^ terminals. No association between aPVT^D2(−)^ terminal responses and return trial order at trial termination (*left*) nor at trigger zone entry (*right*).

**Supplementary Figure 4.**
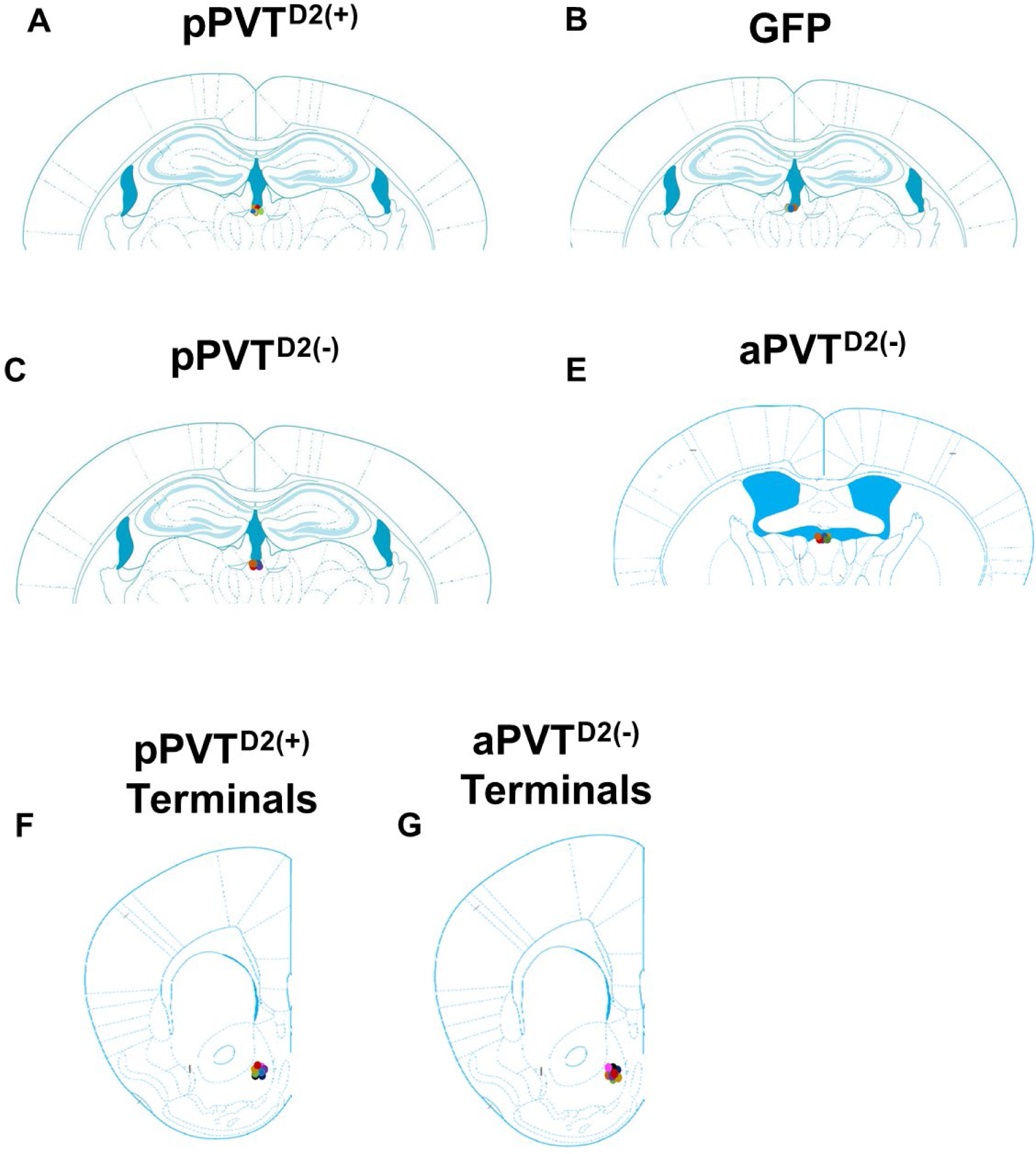
Optical fiber placements. **(A)** Fiber implant location in the pPVT for D2(+) photometry recordings used for experiments in Fig. 1 -pPVT^D2(+)^. **(B)** Fiber implant location in pPVT for GFP controls used in experiments in Fig. 1. **(C)** Fiber implant location in the pPVT for D2(−) photometry recordings used in experiments in Fig. 2 - pPVT^D2(−)^. **(D)** Fiber implant location in the aPVT for D2(−) photometry recordings used for experiments in Fig. 2 - aPVT^D2(−)^. **(E)** NAc fiber implants for pPVT^D2(+)^ terminal photometry recordings used for experiments in Fig. 4A-L. **(F)** NAc fiber implants for aPVT^D2(−)^ terminal photometry recordings used for experiments in Fig. 4M-X.

## METHODS

### Mice

For all experiments, both male and female *Drd2*-Cre mice (8–15 weeks of age) were used. These mice were obtained from the GENSAT (founder line ER44) and were group-housed under a 12-h light-dark cycle (6 a.m. to 6 p.m. light) at a temperature of 70–74 °F and 40–65% humidity. After surgery, mice were single-housed and were provided with food and water *ad-libitum.* Food was restricted for two days before initiating behavioral training procedures, and mice were fed accordingly to maintain 85% of their free-feeding weight. After all testing was completed, mice returned to *ad-libitum* feeding. Animals were randomly allocated to the different experimental conditions reported in this study. Notably, all procedures were performed in accordance with the *Guide for the Care and Use of Laboratory Animals* and were approved by the National Institute of Mental Health (NIMH) Animal Care and Use Committee.

### Viral vectors

AAV9-hSyn-FLEX-GCaMP6s-WPRE-SV40 was produced by the Vector Core of the University of Pennsylvania. AAV9-CAG-Flex-GFP was produced by the University of North Carolina, Chapel Hill (UNC) Vector Core. AAV9-Syn-DO-GCaMP6s was produced by Charu Ramakrishnan (Deisseroth laboratory, Stanford University, CA, USA).

### Stereotaxic surgery

Mice were first anesthetized with a Ketamine/Xylazine solution, and an AngleTwo stereotaxic device (Leica Biosystems) was used for viral injections (approximately one μl) at the following stereotaxic coordinates (at a 6°angle): pPVT, −1.60 mm from Bregma, 0.06 mm lateral from midline, and −3.30 mm vertical from the cortical surface; aPVT, −0.30 mm from bregma, 0.00 mm lateral from the midline and −4.30 mm vertical from the cortical surface. For the fiber photometry experiments, optical fibers with diameters of 400 μm (0.66 NA - Doric Lenses) were used. These fibers were implanted over the pPVT or aPVT immediately following viral injections (targeted 300-400 μm above the injection site). NAc fibers coordinates were: 1.40 mm from bregma, 0.70 mm lateral from the midline, and −4.80 mm vertical from the cortical surface. Fibers were cemented using C&B Metabond Quick Adhesive Cement System (Parkell, Inc.) and Jet Brand dental acrylic (Lang Dental Manufacturing Co., Inc.). For analgesia and anti-inflammatory purposes postoperatively, mice received subcutaneous injections with metacam (meloxicam, 1–2 mg/kg) and were allowed to recover on a heating pad, where they were constantly monitored. Following stereotaxic injections, AAVs were allowed for 2–3 weeks for maximal expression.

### Foraging-like reward-seeking task

The foraging track consisted of a linear maze (150 × 32 × 25, L ×W× H in cm) containing digital distance sensors (15 cm, Pololu Robotics, and Electronics) located throughout the track, which allowed us to track the movement of the mice through the maze. The maze consisted of three zones: trigger zone, corridor, and reward zone. The opposite ends of the track were designated as the trigger zone and reward zone and were connected by a long corridor. Training in the task consisted of two initial sessions of magazine training, in which ¾ of the maze was closed, and mice received 100 µls of the reward every minute for 60 mins. After that, mice were trained in the task for at least 14 days until they reliably performed at least 50 trials or more. The task consisted of 60 min-long sessions of self-paced trials, and mice received one testing session per day. Food-restricted mice were trained to wait in the trigger zone for two seconds for each trial. An auditory cue was then presented, which signaled reward availability, allowing mice to run from the trigger zone, down the corridor, and into the reward zone to retrieve a food reward (strawberry Ensure^®^). Delivery and food consumption signaled trial termination, and therefore, mice had to return to the trigger zone to initiate another trial. Experimental schedule and data acquisition were implemented through the Abet II software for operant control (Lafayette Instruments Neuroscience) and through the Whisker multimedia software (Lafayette Instruments Neuroscience).

### Bulk Ca^2+^ and fiber photometry

All photometry experiments were performed using an RZ5P acquisition system (Tucker-Davis Technologies; TDT) equipped with a real-time signal processor and controlled by a software interface (Synapse version 92). Specifically, the system is integrated with two continuous sinusoidally modulated LEDs (DC4100, ThorLabs) at 473 nm (211 Hz) and 405 nm (531 Hz) that serve as the light source to excite GCaMP6s and an isosbestic autofluorescence signal, respectively. The LED intensity (ranging from 10–15 μW) at the interface between the fiber tip and the animal was constant throughout the session. Fluorescence signals were collected by the same fiber implant that was coupled to a 400 μm optical patch-cord (0.48 NA) and focused onto two separate photoreceivers (2151, Newport Corporation) connected to the emission ports of a custom-ordered fiber photometry Mini Cube (Doric Lenses). TTL pulses recorded by the same system were used to annotate the occurrence of behavioral manipulations. For the measurements of fluorescent calcium signals and ΔF/F analysis, a least-squares linear fit to the 405 nm signal to align it to the 470 nm signal was first applied. The resulting fitted 405 nm signal was then used to normalize the 473 nm as follows: ΔF/F = (473 nm signal − fitted 405 nm signal)/fitted 405 nm signal. All GCaMP6s signal data is presented as the z-score of the dF/F from a baseline prior to the onset of events.

### In-vivo dynamics analyses

Upon calculating the z-score of the dF/F for each event in every trial the mice performed, the latency to the reward zone was calculated. All trials that were higher than 11 seconds and the events corresponding to the trial (cue, approach, reward zone arrival, reward delivery, and consumption) were excluded. Approximately more than 75% of the trials met this inclusion criterion (Supp. Fig. 1F). After that, the trials and events for each individual testing session were sorted by either their latency to approach the reward zone (Approach Latency Blocks) or by trial order - the time in which the trial was completed within a testing session (Trial Group Blocks). After sorting the trials in each testing session, these were then divided into 5 equivalent trial blocks.

Slope-of-the-line calculations: Activity traces for reward approach and reward termination included the activity from time 0, which corresponded to cue delivery, to 11 sec, which was our cut-off for included trials. The maximum and minimum peaks were calculated for each trace, and 80% and 20% values of the max or min peaks were determined. Thereafter, the slope-of-the-line was fitted for the corresponding 80% and 20% values.

### Functional Linear Mixed Model Encoder (FLMM)

The basis for the FLMM analysis was previously described in detail^23^. Briefly, the FLMM analysis is a statistical framework that combines linear mixed models (LMM) and functional regression. The LMM portion of the analysis is a statistical method for data that contains repeated measures and tests the association between independent factors or covariates and dependent factors or outcomes. As such, the LMM allows for the analysis of nested experiments, such as those that contain multiple stimuli and behaviors. The LMM analyses are then combined with functional regression, which takes advantage of autocorrelations in the signal and allows for the calculation of the joint 95% coefficient intervals (CIs) that can be used as the equivalent of other familiar statistical tests (e.g., ANOVAs, correlations). The FLMM framework outputs a plot for each covariate in the model (i.e., the effects of latency and the effects of trial order) and shows whether these are significantly associated with the photometry signal at each time point during the behavior. These FLMM output plots consist of the regression coefficient estimates associated with each covariate at a particular point in the trial (solid lines). The plots also show the estimated covariance function using a pointwise 95% CI for the coefficients of each point (dark-shaded area). Lastly, the plot also shows the calculated joint 95% CI (light-shaded area), which allows for the joint interpretations of each covariate effect on the photometry signals across time intervals at multiple locations in the trial along the functional domain. Statistical significance for the effect of a covariate on the photometry signal can be readily observed in these output plots. As such, the points in the plot where pointwise CI does not contain zero indicate the pointwise statistical significance, and the points where the joint CI does not contain zero indicate jointly statistical significance.

### Histology

To verify GCaMP6s expression and optical fiber placements, mice were injected with euthanasia solution and subsequently sacrificed via transcardial perfusion, first with PBS and then with paraformaldehyde (PFA; 4% in PBS). Brains were then post-fixed in 4% PFA at 4 °C overnight and cryoprotected using a 30% PBS-buffered sucrose solution for ∼24–36 h. Coronal brain sections (45 μm) were generated using a freezing microtome (SM 2010R, Leica). Images were taken using a Carl Zeiss LSM 780 confocal microscope running ZEN software (version 2.3, Carl Zeiss Microscopy, LLC). Optical fiber placements for all subjects included in this study are presented in Supplemental Fig. 4. Mice without correct targeting of optical fibers or GCaMP6s expression were excluded from this study. Correct viral expression for pPVT^D2(+)^ neurons was determined by following the D2R expression reported in our previous studies^4,16^. Correct optical fiber placement was assessed by whether the tip of the fiber was immediately above the pPVT, as shown in Fig. 1A. No animals were excluded from this group. Correct viral expression for PVT^D2(−)^ also followed the D2R(−) expression shown in our previous studies^16^. Particularly for pPVT, any animals that showed virus expression in the medial or lateral habenular or in the ventral medial dorsal thalamus were excluded. For aPVT, any animal that showed virus expression in the paretenial thalamus was excluded from the study. For these animals, correct optical placement was again assessed by whether the tip of the fiber was immediately above pPVT or aPVT, respectively. For the pPVT^D2(−)^ group, one mouse was excluded for incorrect virus expression, and for the aPVT^D2(−)^ group, four mice were excluded for incorrect virus expression and incorrect optical fiber placement.

### Statistics and data presentation

All data were analyzed using GraphPad Prism (Domatics) and Origin Pro 2016 (OriginLab Corp). **The FLMM analysis used R Studio and the ‘FastFLMM’ and ‘ggplot2’ packages.** All statistical tests are indicated when used. No assumptions or corrections were made prior to data analysis. For statistical analyses, a two-tailed paired t-test (nonparametric; test statistic: t) and a repeated measures ANOVA (nonparametric; test statistic: F) were used. ANOVA was followed by a post-hoc Tukey multiple comparisons test if the omnibus test detected a significant difference. All data are presented as mean ± s.e.m. The sample sizes used in our study are similar to those of prior studies^10^. For all the groups tested, the sample size was at least 6 mice, with the exception of the terminal imaging of pPVT^D2(+)^ neurons, for which there were 4 mice. (Supp. Fig. 1H). This was due to technical challenges associated with performing calcium imaging from terminals in our task. All experiments were replicated at least once. The data were assumed to be normal, but this was not formally tested.

### Data availability

All the data that support the findings presented in this study are available from the corresponding author upon reasonable request.

## REFERENCES

1. Zhu, Y., Wienecke, C.F., Nachtrab, G., and Chen, X. (2016). A thalamic input to the nucleus accumbens mediates opiate dependence. Nature 530, 219–222. 10.1038/nature16954.

2. Parsons, M.P., Li, S., and Kirouac, G.J. (2007). Functional and anatomical connection between the paraventricular nucleus of the thalamus and dopamine fibers of the nucleus accumbens. J Comp Neurol 500, 1050–1063. 10.1002/cne.21224.

3. Engelke, D.S., Zhang, X.O., O’Malley, J.J., Fernandez-Leon, J.A., Li, S., Kirouac, G.J., Beierlein, M., and Do-Monte, F.H. (2021). A hypothalamic-thalamostriatal circuit that controls approach-avoidance conflict in rats. Nat Commun 12, 2517. 10.1038/s41467-021-22730-y.

4. Beas, B.S., Wright, B.J., Skirzewski, M., Leng, Y., Hyun, J.H., Koita, O., Ringelberg, N., Kwon, H.B., Buonanno, A., and Penzo, M.A. (2018). The locus coeruleus drives disinhibition in the midline thalamus via a dopaminergic mechanism. Nat Neurosci 21, 963–973. 10.1038/s41593-018-0167-4.

5. Ma, J., du Hoffmann, J., Kindel, M., Beas, B.S., Chudasama, Y., and Penzo, M.A. (2021). Divergent projections of the paraventricular nucleus of the thalamus mediate the selection of passive and active defensive behaviors. Nat Neurosci 24, 1429–1440. 10.1038/s41593-021-00912-7.

6. Do-Monte, F.H., Minier-Toribio, A., Quinones-Laracuente, K., Medina-Colon, E.M., and Quirk, G.J. (2017). Thalamic Regulation of Sucrose Seeking during Unexpected Reward Omission. Neuron 94, 388–400 e384. 10.1016/j.neuron.2017.03.036.

7. Campus, P., Covelo, I.R., Kim, Y., Parsegian, A., Kuhn, B.N., Lopez, S.A., Neumaier, J.F., Ferguson, S.M., Solberg Woods, L.C., Sarter, M., and Flagel, S.B. (2019). The paraventricular thalamus is a critical mediator of top-down control of cue-motivated behavior in rats. Elife 8. 10.7554/eLife.49041.

8. McGinty, J.F., and Otis, J.M. (2020). Heterogeneity in the Paraventricular Thalamus: The Traffic Light of Motivated Behaviors. Front Behav Neurosci 14, 590528. 10.3389/fnbeh.2020.590528.

9. Kirouac, G.J. (2015). Placing the paraventricular nucleus of the thalamus within the brain circuits that control behavior. Neurosci Biobehav Rev 56, 315–329. 10.1016/j.neubiorev.2015.08.005.

10. Otis, J.M., Zhu, M., Namboodiri, V.M.K., Cook, C.A., Kosyk, O., Matan, A.M., Ying, R., Hashikawa, Y., Hashikawa, K., Trujillo-Pisanty, I., et al. (2019). Paraventricular Thalamus Projection Neurons Integrate Cortical and Hypothalamic Signals for Cue-Reward Processing. Neuron 103, 423–431 e424. 10.1016/j.neuron.2019.05.018.

11. Beas, S., Gu, X., Leng, Y., Koita, O., Rodriguez-Gonzalez, S., Kindel, M., Matikainen-Ankney, B.A., Larsen, R.S., Kravitz, A.V., Hoon, M.A., and Penzo, M.A. (2020). A ventrolateral medulla-midline thalamic circuit for hypoglycemic feeding. Nat Commun 11, 6218. 10.1038/s41467-020-19980-7.

12. Iglesias, A.G., and Flagel, S.B. (2021). The Paraventricular Thalamus as a Critical Node of Motivated Behavior via the Hypothalamic-Thalamic-Striatal Circuit. Front Integr Neurosci 15, 706713. 10.3389/fnint.2021.706713.

13. Meffre, J., Sicre, M., Diarra, M., Marchessaux, F., Paleressompoulle, D., and Ambroggi, F. (2019). Orexin in the Posterior Paraventricular Thalamus Mediates Hunger-Related Signals in the Nucleus Accumbens Core. Curr Biol 29, 3298–3306 e3294. 10.1016/j.cub.2019.07.069.

14. Kelley, A.E., Baldo, B.A., and Pratt, W.E. (2005). A proposed hypothalamic-thalamic-striatal axis for the integration of energy balance, arousal, and food reward. J Comp Neurol 493, 72–85. 10.1002/cne.20769.

15. Gao, C., Gohel, C.A., Leng, Y., Ma, J., Goldman, D., Levine, A.J., and Penzo, M.A. (2023). Molecular and spatial profiling of the paraventricular nucleus of the thalamus. Elife 12. 10.7554/eLife.81818.

16. Gao, C., Leng, Y., Ma, J., Rooke, V., Rodriguez-Gonzalez, S., Ramakrishnan, C., Deisseroth, K., and Penzo, M.A. (2020). Two genetically, anatomically and functionally distinct cell types segregate across anteroposterior axis of paraventricular thalamus. Nat Neurosci 23, 217–228. 10.1038/s41593-019-0572-3.

17. Averbeck, B.B., and Costa, V.D. (2017). Motivational neural circuits underlying reinforcement learning. Nat Neurosci 20, 505–512. 10.1038/nn.4506.

18. Kelley, A.E. (1999). Neural integrative activities of nucleus accumbens subregions in relation to learning and motivation. Psychobiology 27, 198–213. 10.3758/BF03332114.

19. Shima, Y., Skibbe, H., Sasagawa, Y., Fujimori, N., Iwayama, Y., Isomura-Matoba, A., Yano, M., Ichikawa, T., Nikaido, I., Hattori, N., and Kato, T. (2023). Distinctiveness and continuity in transcriptome and connectivity in the anterior-posterior axis of the paraventricular nucleus of the thalamus. Cell Rep 42, 113309. 10.1016/j.celrep.2023.113309.

20. Kvitsiani, D., Ranade, S., Hangya, B., Taniguchi, H., Huang, J.Z., and Kepecs, A. (2013). Distinct behavioural and network correlates of two interneuron types in prefrontal cortex. Nature 498, 363–366. 10.1038/nature12176.

21. Jensen, T.L., Kiersgaard, M.K., Sorensen, D.B., and Mikkelsen, L.F. (2013). Fasting of mice: a review. Lab Anim 47, 225–240. 10.1177/0023677213501659.

22. Labouebe, G., Boutrel, B., Tarussio, D., and Thorens, B. (2016). Glucose-responsive neurons of the paraventricular thalamus control sucrose-seeking behavior. Nat Neurosci 19, 999–1002. 10.1038/nn.4331.

23. Loewinger, G., Cui, E., Lovinger, D., and Pereira, F. (2023). A Statistical Framework for Analysis of Trial-Level Temporal Dynamics in Fiber Photometry Experiments. bioRxiv. 10.1101/2023.11.06.565896.

24. Cui, E., Leroux, A., Smirnova, E., and Crainiceanu, C.M. (2022). Fast Univariate Inference for Longitudinal Functional Models. J Comput Graph Stat 31, 219–230. 10.1080/10618600.2021.1950006.

25. Zhang, J., Chen, D., Sweeney, P., and Yang, Y. (2020). An excitatory ventromedial hypothalamus to paraventricular thalamus circuit that suppresses food intake. Nat Commun 11, 6326. 10.1038/s41467-020-20093-4.

26. Cheng, J., Wang, J., Ma, X., Ullah, R., Shen, Y., and Zhou, Y.D. (2018). Anterior Paraventricular Thalamus to Nucleus Accumbens Projection Is Involved in Feeding Behavior in a Novel Environment. Front Mol Neurosci 11, 202. 10.3389/fnmol.2018.00202.

27. Millan, E.Z., Ong, Z., and McNally, G.P. (2017). Paraventricular thalamus: Gateway to feeding, appetitive motivation, and drug addiction. Prog Brain Res 235, 113–137. 10.1016/bs.pbr.2017.07.006.

28. Vollmer, K.M., Green, L.M., Grant, R.I., Winston, K.T., Doncheck, E.M., Bowen, C.W., Paniccia, J.E., Clarke, R.E., Tiller, A., Siegler, P.N., et al. (2022). An opioid-gated thalamoaccumbal circuit for the suppression of reward seeking in mice. Nat Commun 13, 6865. 10.1038/s41467-022-34517-w.

29. Lafferty, C.K., Yang, A.K., Mendoza, J.A., and Britt, J.P. (2020). Nucleus Accumbens Cell Type-and Input-Specific Suppression of Unproductive Reward Seeking. Cell Rep 30, 3729–3742 e3723. 10.1016/j.celrep.2020.02.095.

30. Dong, X., Li, S., and Kirouac, G.J. (2017). Collateralization of projections from the paraventricular nucleus of the thalamus to the nucleus accumbens, bed nucleus of the stria terminalis, and central nucleus of the amygdala. Brain Struct Funct 222, 3927–3943. 10.1007/s00429-017-1445-8.

31. Clark, A.M., Leroy, F., Martyniuk, K.M., Feng, W., McManus, E., Bailey, M.R., Javitch, J.A., Balsam, P.D., and Kellendonk, C. (2017). Dopamine D2 Receptors in the Paraventricular Thalamus Attenuate Cocaine Locomotor Sensitization. eNeuro 4. 10.1523/ENEURO.0227-17.2017.

32. Floresco, S.B. (2015). The nucleus accumbens: an interface between cognition, emotion, and action. Annu Rev Psychol 66, 25–52. 10.1146/annurev-psych-010213-115159.

33. Floresco, S.B., McLaughlin, R.J., and Haluk, D.M. (2008). Opposing roles for the nucleus accumbens core and shell in cue-induced reinstatement of food-seeking behavior. Neuroscience 154, 877–884. 10.1016/j.neuroscience.2008.04.004.

34. Ikemoto, S., and Panksepp, J. (1999). The role of nucleus accumbens dopamine in motivated behavior: a unifying interpretation with special reference to reward-seeking. Brain Res Brain Res Rev 31, 6–41. 10.1016/s0165-0173(99)00023-5.

35. Zhang, X., and van den Pol, A.N. (2017). Rapid binge-like eating and body weight gain driven by zona incerta GABA neuron activation. Science 356, 853–859. 10.1126/science.aam7100.

36. Ong, Z.Y., Liu, J.J., Pang, Z.P., and Grill, H.J. (2017). Paraventricular Thalamic Control of Food Intake and Reward: Role of Glucagon-Like Peptide-1 Receptor Signaling. Neuropsychopharmacology 42, 2387–2397. 10.1038/npp.2017.150.

37. Ye, Q., Nunez, J., and Zhang, X. (2022). Oxytocin Receptor-Expressing Neurons in the Paraventricular Thalamus Regulate Feeding Motivation through Excitatory Projections to the Nucleus Accumbens Core. J Neurosci 42, 3949–3964. 10.1523/JNEUROSCI.2042-21.2022.

38. Betley, J.N., Cao, Z.F., Ritola, K.D., and Sternson, S.M. (2013). Parallel, redundant circuit organization for homeostatic control of feeding behavior. Cell 155, 1337–1350. 10.1016/j.cell.2013.11.002.

39. Penzo, M.A., and Gao, C. (2021). The paraventricular nucleus of the thalamus: an integrative node underlying homeostatic behavior. Trends Neurosci 44, 538–549. 10.1016/j.tins.2021.03.001.

40. Horio, N., and Liberles, S.D. (2021). Hunger enhances food-odour attraction through a neuropeptide Y spotlight. Nature 592, 262–266. 10.1038/s41586-021-03299-4.

41. Dong, X., Li, S., and Kirouac, G.J. (2020). A projection from the paraventricular nucleus of the thalamus to the shell of the nucleus accumbens contributes to footshock stress-induced social avoidance. Neurobiol Stress 13, 100266. 10.1016/j.ynstr.2020.100266.

42. Paniccia, J.E., Vollmer, K.M., Green, L.M., Grant, R.I., Winston, K.T., Buchmaier, S., Westphal, A.M., Clarke, R.E., Doncheck, E.M., Bordieanu, B., et al. (2023). Restoration of a paraventricular thalamo-accumbal behavioral suppression circuit prevents reinstatement of heroin seeking. Neuron. 10.1016/j.neuron.2023.11.024.

43. Al-Hasani, R., McCall, J.G., Shin, G., Gomez, A.M., Schmitz, G.P., Bernardi, J.M., Pyo, C.O., Park, S.I., Marcinkiewcz, C.M., Crowley, N.A., et al. (2015). Distinct Subpopulations of Nucleus Accumbens Dynorphin Neurons Drive Aversion and Reward. Neuron 87, 1063–1077. 10.1016/j.neuron.2015.08.019.

44. Chen, G., Lai, S., Bao, G., Ke, J., Meng, X., Lu, S., Wu, X., Xu, H., Wu, F., Xu, Y., et al. (2023). Distinct reward processing by subregions of the nucleus accumbens. Cell Rep 42, 112069. 10.1016/j.celrep.2023.112069.

45. Lafferty, C.K., and Britt, J.P. (2020). Off-Target Influences of Arch-Mediated Axon Terminal Inhibition on Network Activity and Behavior. Front Neural Circuits 14, 10. 10.3389/fncir.2020.00010.

46. Baimel, C., Jang, E., Scudder, S.L., Manoocheri, K., and Carter, A.G. (2022). Hippocampal-evoked inhibition of cholinergic interneurons in the nucleus accumbens. Cell Rep 40, 111042. 10.1016/j.celrep.2022.111042.

47. Giannotti, G., Gong, S., Fayette, N., Heinsbroek, J.A., Orfila, J.E., Herson, P.S., Ford, C.P., and Peters, J. (2021). Extinction blunts paraventricular thalamic contributions to heroin relapse. Cell Rep 36, 109605. 10.1016/j.celrep.2021.109605.

48. Joffe, M.E., and Grueter, B.A. (2016). Cocaine Experience Enhances Thalamo-Accumbens N-Methyl-D-Aspartate Receptor Function. Biol Psychiatry 80, 671–681. 10.1016/j.biopsych.2016.04.002.

49. Arcelli, P., Frassoni, C., Regondi, M.C., De Biasi, S., and Spreafico, R. (1997). GABAergic neurons in mammalian thalamus: a marker of thalamic complexity? Brain Res Bull 42, 27–37. 10.1016/s0361-9230(96)00107-4.

50. Halassa, M.M., and Acsady, L. (2016). Thalamic Inhibition: Diverse Sources, Diverse Scales. Trends Neurosci 39, 680–693. 10.1016/j.tins.2016.08.001.

51. Halassa, M.M., Chen, Z., Wimmer, R.D., Brunetti, P.M., Zhao, S., Zikopoulos, B., Wang, F., Brown, E.N., and Wilson, M.A. (2014). State-dependent architecture of thalamic reticular subnetworks. Cell 158, 808–821. 10.1016/j.cell.2014.06.025.

52. Wimmer, R.D., Schmitt, L.I., Davidson, T.J., Nakajima, M., Deisseroth, K., and Halassa, M.M. (2015). Thalamic control of sensory selection in divided attention. Nature 526, 705–709. 10.1038/nature15398.

53. Zikopoulos, B., and Barbas, H. (2007). Circuits formultisensory integration and attentional modulation through the prefrontal cortex and the thalamic reticular nucleus in primates. Rev Neurosci 18, 417–438. 10.1515/revneuro.2007.18.6.417.

54. Guillery, R.W., and Harting, J.K. (2003). Structure and connections of the thalamic reticular nucleus: Advancing views over half a century. J Comp Neurol 463, 360–371. 10.1002/cne.10738.

55. Brown, J.W., Taheri, A., Kenyon, R.V., Berger-Wolf, T.Y., and Llano, D.A. (2020). Signal Propagation via Open-Loop Intrathalamic Architectures: A Computational Model. eNeuro 7. 10.1523/ENEURO.0441-19.2020.

56. Willis, A.M., Slater, B.J., Gribkova, E.D., and Llano, D.A. (2015). Open-loop organization of thalamic reticular nucleus and dorsal thalamus: a computational model. J Neurophysiol 114, 2353–2367. 10.1152/jn.00926.2014.

57. Quinones-Laracuente, K., Vega-Medina, A., and Quirk, G.J. (2021). Time-Dependent Recruitment of Prelimbic Prefrontal Circuits for Retrieval of Fear Memory. Front Behav Neurosci 15, 665116. 10.3389/fnbeh.2021.665116.

58. Kolaj, M., Zhang, L., Ronnekleiv, O.K., and Renaud, L.P. (2012). Midline thalamic paraventricular nucleus neurons display diurnal variation in resting membrane potentials, conductances, and firing patterns in vitro. J Neurophysiol 107, 1835–1844. 10.1152/jn.00974.2011.

59. Grahek, I., Shenhav, A., Musslick, S., Krebs, R.M., and Koster, E.H.W. (2019). Motivation and cognitive control in depression. Neurosci Biobehav Rev 102, 371–381. 10.1016/j.neubiorev.2019.04.011.

60. Der-Avakian, A., Barnes, S.A., Markou, A., and Pizzagalli, D.A. (2016). Translational Assessment of Reward and Motivational Deficits in Psychiatric Disorders. Curr Top Behav Neurosci 28, 231–262. 10.1007/7854_2015_5004.

